# Sae2/CtIP prevents R-loop accumulation in eukaryotic cells

**DOI:** 10.1101/394924

**Authors:** Nodar Makharashvili, Sucheta Arora, Yizhi Yin, Qiong Fu, Justin W. C. Leung, Kyle M. Miller, Tanya T. Paull

## Abstract

The Sae2/CtIP protein is required for efficient processing of DNA double-strand breaks that initiate homologous recombination in eukaryotic cells. Sae2/CtIP is also important for survival of single-stranded Top1-induced lesions and CtIP is known to associate directly with transcription-associated complexes in mammalian cells. Here we investigate the role of Sae2/CtIP at single-strand lesions in budding yeast and in human cells and find that depletion of Sae2/CtIP promotes the accumulation of stalled RNA polymerase and RNA-DNA hybrids at sites of highly expressed genes. Overexpression of the RNA-DNA helicase Senataxin suppresses DNA damage sensitivity and R-loop accumulation in Sae2/CtIP-deficient cells, and a catalytic mutant of CtIP fails to complement this sensitivity, indicating a role for CtIP nuclease activity in the repair process. Based on this evidence, we propose that R-loop processing by 5’ flap endonucleases is a necessary step in the stabilization and removal of nascent R-loop initiating structures in eukaryotic cells.

## Introduction

Double-strand breaks in DNA are known to be lethal lesions in eukaryotic cells, and can be an important source of genomic instability during oncogenic transformation because of the possibility of misrepair, translocations, and rearrangements that initiate from these lesions ^1^. The repair of double-strand breaks in eukaryotes occurs through pathways related to either non-homologous end joining or homologous recombination, although many variations on these basic pathways can occur in cells depending on the cell cycle phase, the extent of DNA end processing that occurs, which enzymes are utilized to do the processing, and what resolution outcomes predominate ^2–4^.

The Sae2/CtIP enzyme is important for DNA end processing in eukaryotes and has been shown to act in several ways to facilitate the removal of the 5’ strand at DNA double-strand breaks ^3^. Phosphorylated Sae2/CtIP promotes the activity of the Mre11 nuclease in the Mre11/Rad50/Xrs2(Nbs1)(MRX(N)) complex, which initiates the trimming of the DNA end on the 5’ strand ^5^. Sae2/CtIP and MRX(N) promote the removal of the Ku heteodimer, which acts as a block to resection during non-homologous end joining ^5–7^ and also recruit the long-range 5’ to 3’ nucleases Exo1 and Dna2 which do extensive processing of the ends ^8–12^.

In addition to the activities of Sae2/CtIP that promote MRX(N) functions, the protein has also been shown to possess intrinsic nuclease activity that is important for the processing of breaks, particularly those formed in the context of protein lesions, radiation-induced DNA damage, or camptothecin (CPT) damage during S phase ^3,13–15^. Nuclease-deficient CtIP fails to complement human cells deficient in CtIP for survival of radiation-induced DNA damage while homologous recombination at restriction enzyme-induced break sites is comparable to wild-type-complemented cells ^14,15^.

Eukaryotic cells lacking Sae2 or its mammalian ortholog CtIP were also observed years ago to be hypersensitive to topoisomerase 1 poisons such as CPT ^16,17^. Top1 bound to CPT is stalled in its catalytic cycle in a covalent tyrosine 3’ linkage with DNA, creating a protein-linked DNA strand adjacent to a 5’ nick ^18^. This lesion targets one DNA strand but can lead to double-strand breaks during replication. Importantly, Top1 is highly active at sites of ongoing transcription due to the need for the release of topological stress in front of and behind the RNA polymerase ^19–21^. Considering that MRX(N) complexes as well as Sae2/CtIP are essential for the removal of 5’ Spo11 conjugates during meiosis ^22–24^ coincident with their role in 5’ strand processing, it is surprising that Sae2/CtIP-deficient cells exhibit such sensitivity to 3’ single-strand lesions, and the mechanistic role that Sae2/CtIP plays at these lesions is currently unknown.

In recent years, it has become clear that transcription can play a major role in promoting genomic instability by forming stalled transcription complexes and RNA-DNA hybrids in the genome ^25^. Stable annealing of nascent RNA with the DNA template strand can occur at sites of stalled RNA polymerase complexes, and the “R-loops” formed in this way can block replication as well as other DNA transactions ^26^. A wealth of evidence accumulated recently suggests that these events can lead to single-strand and double-strand breaks in DNA that provide recombinogenic intermediates for misrepair events ^27,28^.

The Sen1 protein in budding yeast has been shown to regulate many aspects of RNA biology, including termination of RNA polymerase II transcription, 3′ end processing of mRNA, and dissociation of RNA-DNA hybrids ^29–31^. The human ortholog of Sen1, Senataxin, has also been shown to resolve RNA-DNA hybrids as well as to associate with replication forks to protect fork integrity when traversing transcribed genes ^32–35^. Mutations in the gene encoding Senataxin are responsible to the neurodegenerative disorder Ataxia with Oculomotor Apraxia 2 as well as an early-onset form of amyotrophic lateral sclerosis^36^.

In this work we sought to understand the mechanistic basis of the hypersensitivity of Sae2/CtIP-deficient cells to Top1 poisons, finding unexpectedly that this is dependent on active transcription. Genetic evidence in yeast and human cells shows that Sae2/CtIP-deficient cells require the RNA-DNA helicase Senataxin for survival of genotoxic agents and that these cells exhibit high levels of RNA polymerase stalling and R-loop formation, consistent with a failure to recognize or process RNA-DNA hybrids. Lastly, we provide evidence for an important role of Sae2/CtIP in processing R-loops that promotes the action of RNA-DNA helicases and ultimately cell survival after DNA damage.

## Results

### Transcription termination factors rescue DNA damage sensitivity of Δsae2 and mre11 nuclease-deficient yeast cells

To test for an effect of transcriptional regulation on the *Δsae2* phenotype in yeast, we overexpressed several different RNA Pol II-associated factors in the mutant strain. We found that overexpression of the termination factor SEN1 markedly improved survival of the *Δsae2* strain to genotoxic agents (Fig. 1A). *S. cerevisiae SEN1* encodes a helicase that is responsible for unwinding RNA-DNA hybrids and also promotes transcription termination through direct contact with RNA Pol II as well as 3′ end processing of RNA ^37^. The ability of SEN1 overexpression to partially alleviate the toxicity of CPT was also observed with the Mre11 nuclease-deficient mutant *mre11- H125N* ^38^ and particularly with the double mutant *Δsae2 mre11-H125N* (Fig. 1B). A mutation located in the conserved helicase domain of SEN1 (G1747D) reduces the ability of SEN1 to overcome CPT toxicity in the *Δsae2* strain (Fig. 1A) but there was no effect of R302W, a mutation reported to block binding to the Rpb1 subunit of RNA Pol II ^39^. The *sen1-G1747D* mutant is deficient in transcription termination but not in 3′ end processing of RNA ^30^, thus we conclude that the termination function of the Sen1 enzyme is important for the rescue of CPT sensitivity in *Δsae2* strains. In contrast, SEN1 overexpression in *Δsae2* has no effect on the efficiency of resection (Fig. S1), as measured in an assay for single-strand annealing ^40^ previously shown be dependent on *SAE2* due to its importance in DNA end resection ^41^.

**Figure 1.**
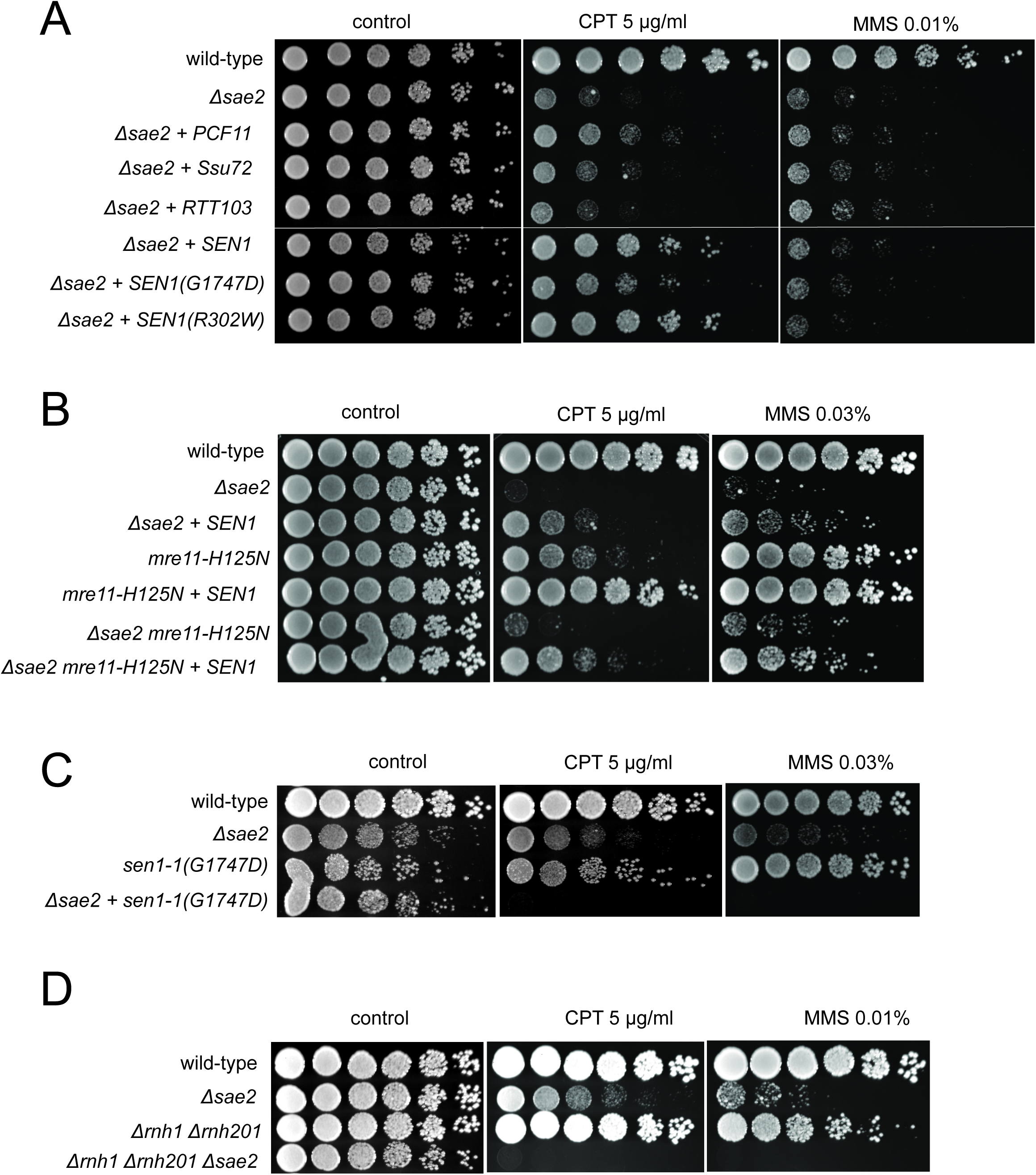
Transcription termination factors suppress DNA damage sensitivity of *sae2* and *mre11* nuclease-deficient strains. (A) Full-length *PCF11, SSU72, RTT103, SEN1*, and *sen1* mutants G1747D and R302W were expressed from a 2μ plasmid in *Δsae2* cells. Fivefold serial dilutions of cells expressing the indicated Sae2 alleles were plated on nonselective media (control) or media containing camptothecin (CPT, 5.0 μg/ml) or methylmethanesulfonate (MMS, 0.01%) and grown for 48 hrs (control) or 70 hrs (CPT and MMS). (B) *SEN1* was expressed from a 2μ plasmid in *Δsae2, mre11- H125N, and Δsae2 mre11-H125N* cells and analyzed for CPT sensitivity as in (A). (C) Wild-type, *Δsae2, sen1-1(G1747D)*, and *Δsae2 sen1-1(G1747D)* strains were analyzed as in (A). (D) Wild-type, *Δsae2, δrnh1 δrnh201*, and *Δsae2 δrnh1 δrnh201* strains were analyzed as in (A).

To further investigate the genetic relationship between *SEN1* and *Δsae2* phenotypes, we deleted the *SAE2* gene in a *sen1-1* (G1747D) background. A complete deletion of *SEN1* is lethal ^42^; however, the *sen1-1* allele has been used as a hypomorphic mutant and is deficient in transcription termination and removal of R-loops in vivo ^30^. Although the *sen1-1* strain is not sensitive to the levels of DNA damaging agents used here, a combination with *Δsae2* generates extreme sensitivity to both CPT and MMS (Fig. 1C). Synthetic sensitivity of *sen1-1* with other DNA repair mutant strains has previously been shown for HU exposure ^30^. Since the Sen1 helicase acts to remove R-loops from genomic loci, we also tested whether *Δsae2* strains show synthetic sensitivity to CPT in combination with deletions of RNase H enzymes which remove ribonucleotides from DNA. Deletion of both RNase H1 and H2 in a *δrnh1 δrnh201* strain generates DNA damage sensitivity as previously shown ^43–45^; however, this is further exacerbated by a deletion of *SAE2* (Fig. 1D). Taken together, these results suggest that loss of Sae2, either by itself or in combination with Mre11 nuclease activity, generates a form of DNA damage sensitivity that requires efficient removal of RNA or ribonucleotides from DNA.

PCF11 is a component of the cleavage and polyadenylation complex (CPAC) which promotes termination of transcription by direct interaction with phosphorylated RNA Pol II as well as promoting the cleavage and processing of nascent mRNA ^46,47^. Here we found that PCF11 also improved the survival of yeast strains lacking *SAE2* when tested for survival of CPT but there was little effect of overexpressing other proteins that also regulate transcription through RNA Pol II including *SSU72, RTT103, NRD1*, and *YSH1* (Fig. 1A and data not shown).

RNA-DNA hybrids form at sites of RNA polymerase pausing and at locations of collisions between the transcription machinery and the replication fork ^26^. We postulated that the source of lethal damage in a *Δsae2* strain may be related to transcription-replication conflicts, based on the *SEN1* and RNase H observations above. To test this idea we synchronized wild-type and *Δsae2* yeast strains with alpha factor and exposed the cells to CPT in either G_1_ or S phases of the cell cycle. As expected, the *Δsae2* strain showed marked sensitivity to CPT and this was specific to exposure in S phase (Fig. 2A), suggesting that movement of replication forks through Top1 DNA damage sites is important for the DNA damage sensitivity. We also tested for the effect of transcription on CPT survival by incubating cells with the general transcription inhibitor thiolutin ^48^. Remarkably, inhibition of transcription completely alleviated the sensitivity of the *Δsae2* strain to CPT, generating wild-type levels of survival (Fig. 2B).

**Figure 2.**
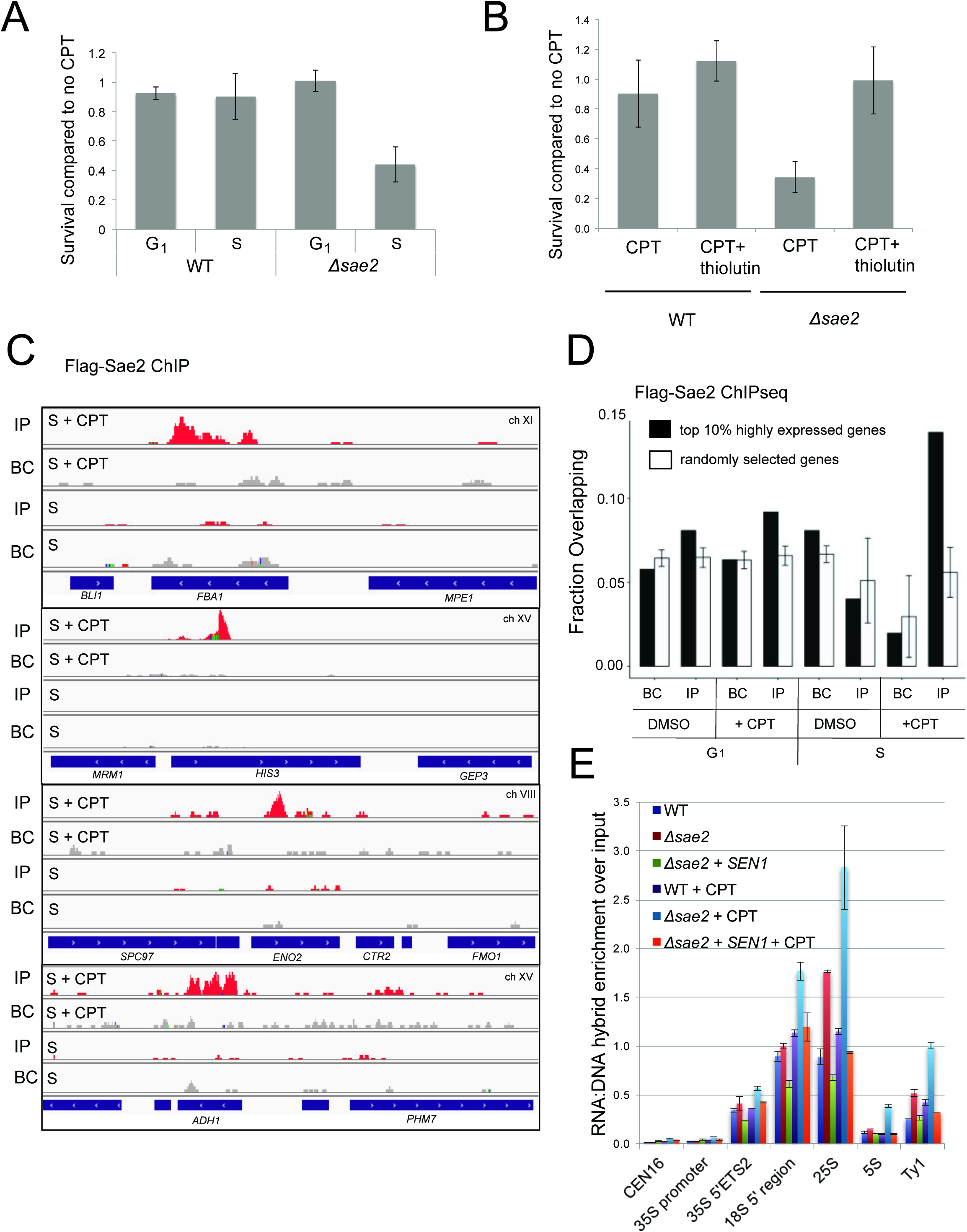
Sae2 associates with sites of high levels of transcription. (A) The survival of wild-type and *Δsae2* strains was measured by exposing cells to camptothecin (100 μM for 2 hrs) while in G_1_ phase or S phase and plating cells on rich media. The percentage of viable colonies is shown relative to cells exposed to DMSO only, with 3 biological replicates (error bars represent standard deviation). (B) The survival of wild-type and *Δsae2* strains was measured in the absence or presence of active transcription by exposing S phase cells to thiolutin (2.5 μg/ml for 30 min) or DMSO prior to camptothecin exposure in S phase as in (A). (C) Representative examples of Sae2-ChIP at the FBA1, HIS3, ENO2, and ADH1 genes in *Δsae2* cells expressing Flag-Sae2, in S phase with CPT exposure as in (A). Reads from the immunoprecipitated sample are shown (IP) in comparison to control immunoprecipitations performed in the absence of Flag antibody (bead control, BC). (D) Data from Sae2 ChIP-seq was compared to previous data on transcription levels in wild-type yeast cells ^50–52^(See Table S2). The overlap between peaks identified by Sae2 ChIP-seq were compared to the top 10% of transcribed genes (486 genes; excluding rDNA loci) or a randomly chosen set of genes. The randomized set comparison was performed 100 times. Error bars represent S. D. (E) R-loops were quantified at various loci using S9.6 immunoprecipitation in wild-type, *Δsae2*, and *Δsae2 +* SEN1 yeast strains, with or without CPT in S phase, as indicated. Levels of DNA sites enriched in S9.6 immunoprecipitations were measured within the rDNA locus, at a control site (CEN16), and at a transposable element, Ty1. Error bars represent standard deviation from 2 biological replicates.

### Sae2 occupancy is elevated at sites of high transcription in S phase cells

To determine if the relationship between *SAE2* deletion and transcription is direct, we sought to determine the genomic locations of Sae2 protein binding in yeast. We performed ChIP assays using Flag-tagged Sae2 and analyzed the peaks in relation to the bead control (no antibody). This analysis revealed small peaks of Sae2 enrichment, primarily in the S phase + CPT sample (176 peaks identified by Model-based Analysis of ChIP-Seq v.2 (MACS2) ^49^ after removal of overlaps with bead control) (Fig. 2C, Table S2). In contrast, only 45 peaks were identified by this criteria in S phase in the absence of DNA damage. The locations of the sites enriched with CPT treatment did not correlate with sites of replication origins (data not shown) but were enriched for highly transcribed genes, measuring the overlap between the binding sites and the top 10% of highly transcribed genes ^50–52^ in comparison to a randomly selected subset (Fig. 2D). The enrichment for Sae2 occupancy at sites of high transcription was only observed with cells in S phase, not with the G_1_ phase cells, and only in cells treated with CPT.

### Δsae2 cells accumulate R-loops and stalled RNA Pol II during CPT exposure

The results from these experiments suggested an involvement of transcription in the DNA damage sensitivity of *Δsae2* cells, possibly related to an accumulation of R-loops. To test this hypothesis we used the S9.6 antibody specific for RNA-DNA hybrids ^53^ in chromatin immunoprecipitation experiments comparing wild-type, *Δsae2*, and *Δsae2* with SEN1 overexpression. Unfortunately our S9.6 immunoprecipitation experiments in yeast did not yield high enough signals for analysis of Pol II transcribed genes, but we did find that CPT exposure induces R-loops throughout the rDNA locus (Fig. 2E). At several sites in this region, *Δsae2* cells exposed to CPT showed an enrichment of R-loops, particularly in the 18S and 25S rDNA genes.

If Sae2 is localized to sites of high transcription and its loss is partially alleviated by enzymes that promote transcription termination, we reasoned that levels of RNA polymerase may be stalled at these sites in *Δsae2* strains. To test this idea we utilized a tagged RNA Pol II strain (HTB-Rpb2) ^54^ to monitor the occupancy of RNA Pol II at sites in the genome with high levels of constitutive transcription where Sae2 was observed by ChIP-qPCR in Fig. 2C. HTB-tagged Pol II complexes were isolated from wild-type, *Δsae2*, and *Δsae2* with *SEN1* overexpression strains that were synchronized in G_1_ with alpha-factor and released into S phase. All of the strains showed very similar levels of RNA Pol II occupancy on the *ADH1, ENO2*, and *FBA1* genes in the absence of DNA damage (Fig. 3A). In contrast, when the strains were released into S phase in the presence of CPT, the wild-type strain showed an average of 1.5 to 2-fold higher levels of RNA Pol II occupancy, while *Δsae2* strains exhibited 2.5 to 5.5-fold higher levels of polymerase pausing (Fig. 3A). Importantly, *Δsae2* with SEN1 overexpression showed reduced levels of polymerase occupancy, similar to the wild-type strain with CPT.

**Figure 3.**
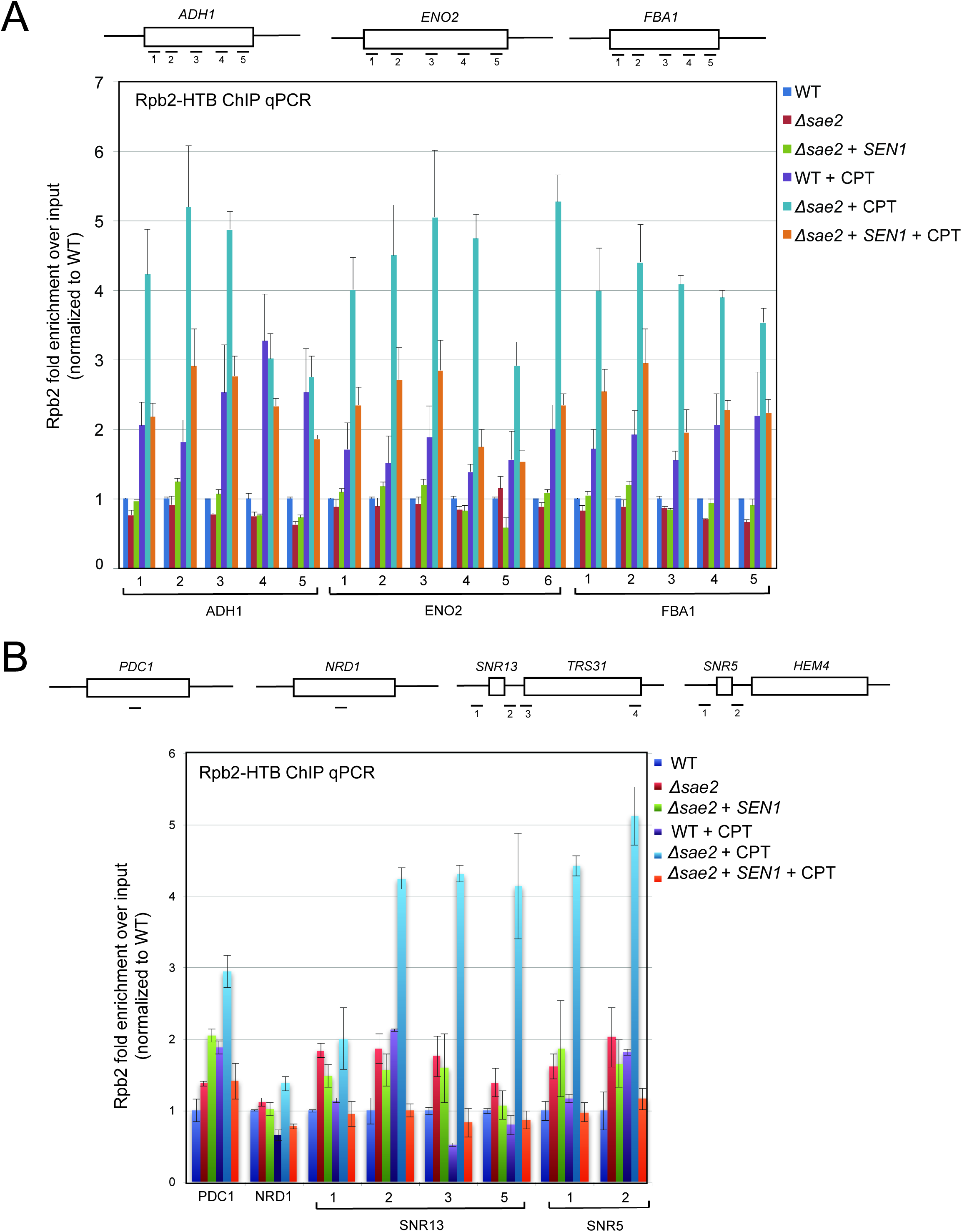
RNA Polymerase stalling at highly transcribed genes is exacerbated in *sae2* strains with DNA damage. (A) HTB-tagged Rpb2 (a component of RNA Pol II) levels were measured at various genes during S phase in wild-type and *Δsae2* strains with or without CPT exposure and in the presence or absence of overexpressed SEN1, as indicated. Enrichment relative to input DNA is shown, with all values normalized to values obtained with the wild-type strain in the absence of damage. Error bars represent standard deviation from 2 biological replicates. Approximate locations of primer sets relative to the gene are shown. (B) HTB-tagged Rpb2 levels were measured at various genes as in (A). *SNR5* and *SNR13* are non-coding nucleolar RNAs.

We also examined RNA Pol II occupancy at other genomic locations where transcription termination has previously been shown to be regulated by Sen1. We did not observe any CPT-induced RNA Pol II pausing at *PDC1* or *NRD1* genes, both known to be dependent on SEN1 ^32,47^ but not identified in our Sae2 ChIP experiment. However, pausing in the absence of Sae2 was induced by CPT at the snoRNA genes *SNR13* and *SNR5* (Fig. 3B), which have been shown to be transcribed by RNA Pol II and exhibit termination read-through, altered RNA Pol II occupancy, and R-loops in the absence of *SEN1* function ^31,47^. At both of these genes and the *TRS31* gene adjacent to *SNR13* we found high levels of RNA Pol II pausing with CPT treatment in the *Δsae2* strain, which was alleviated by *SEN1* overexpression. Overall, these results are consistent with the hypothesis that Sae2 is present at a subset of highly transcribed genes during CPT exposure, and that high levels of toxic R-loops form at these sites in *Δsae2* strains which can be reduced by *SEN1*.

### The CPT sensitivity of CtIP deficient cells is rescued by over-expression of Senataxin or inhibition of transcription

The functional ortholog of *SAE2* in mammalian cells is CtIP ^17^, which promotes resection of DNA double-strand breaks in conjunction with the MRN complex ^55^. It was previously shown that depletion of CtIP generates extreme sensitivity to topoisomerase poison induced DNA lesions, particularly CPT-induced DNA damage ^17,55–57^. To examine the role of CtIP in human cells we used a U2OS cell line with a stably integrated doxycycline-inducible CtIP shRNA cassette. The cells were complemented with either shRNA-resistant wild-type eGFP-CtIP, or with vector only (Fig. S2). As expected, depletion of CtIP greatly diminishes cell survival in the presence of CPT, and re-expression of wild-type CtIP rescues the CPT sensitivity (Fig. 4A).

**Figure 4.**
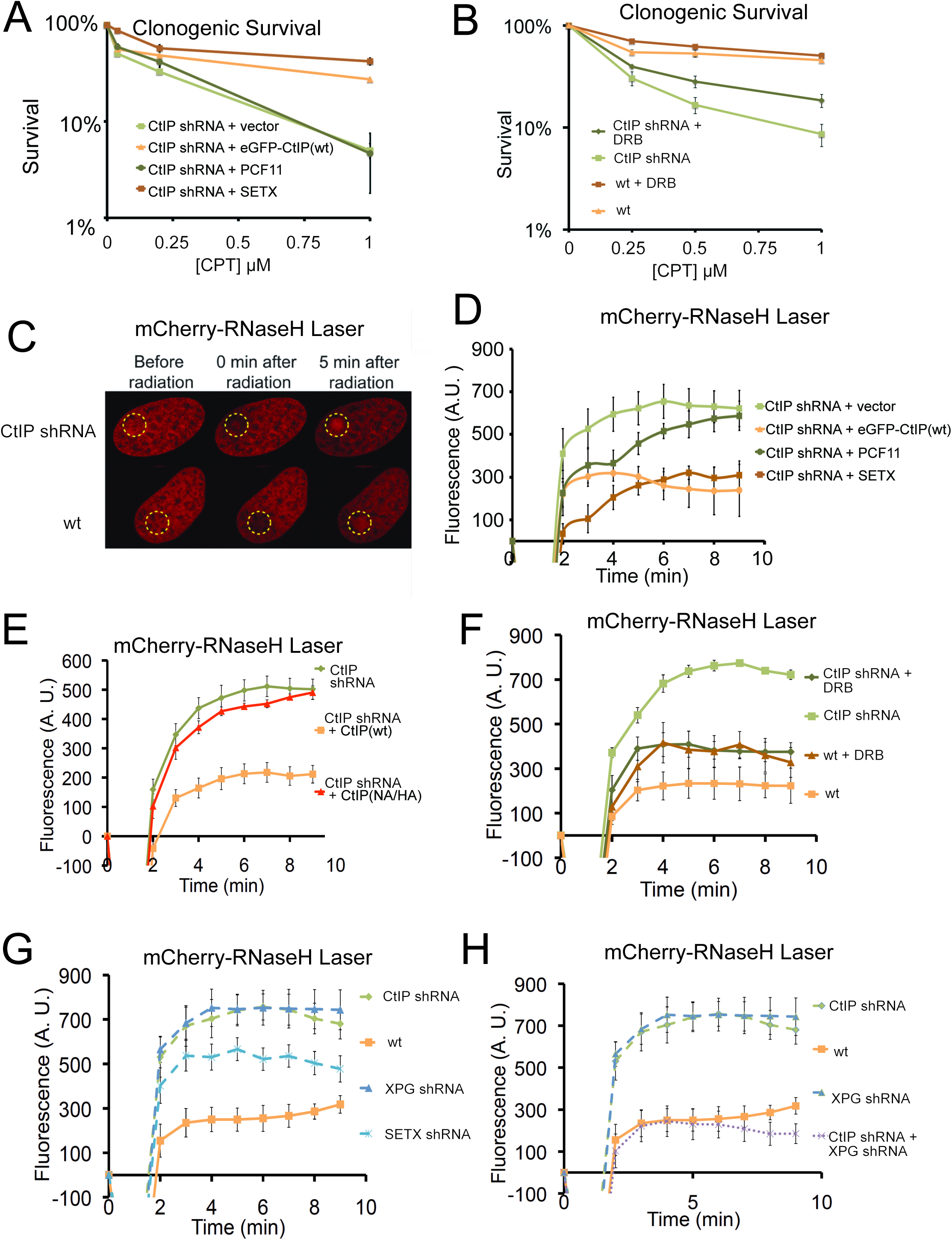
CtIP deficiency induces R-loop accumulation at sites of DNA damage. (A) CtIP-depleted U2OS cells complemented with vector only or constructs overexpressing eGFP-CtIP, PCF11, or Senataxin (C-terminus) as indicated were exposed to increasing concentrations of CPT (1 hr). Cell viability was determined by clonogenic survival assay in comparison to untreated cells. Results are shown from 3 biological replicates and error bars represent S. D. (B) Wild-type or CtIP-depleted U2OS cells were pre-treated with DRB (20 μM) prior to CPT exposure and cell viability was determined as in (A). (C) Live cell imaging was performed with U2OS cells stably expressing RNaseH^D10R-E48R^-mCherry. The circle indicates the site of laser damage. (D) CtIP-depleted U2OS cells were complemented with eGFP-CtIP, PCF11, or Senataxin (C-terminus) as indicated and RNaseH^D10R-E48R^-mCherry accumulation at the laser damage was quantified. The average of 4 cells is shown and error bars represent S. E. M. (E) RNaseH^D10R-E48R^-mCherry accumulation at laser damage sites was measured in CtIP-depleted U2OS cells complemented with wild-type or nuclease deficient (“NA/HA”) eGFP-CtIP as in (D); n > 12, error bars represent S.E.M. (F) Wild-type or CtIP-depleted U2OS cells were pre-treated with DRB (20 μM) prior to measurement of RNaseH^D10R-E48R^-mCherry accumulation at laser damage sites as in (D); n > 5; error bars represent S.E.M. (G) RNaseH^D10R-E48R^-mCherry accumulation at laser damage sites was measured in wild-type or CtIP-depleted, XPG-depleted, or Senataxin (SETX)-depleted U2OS cells as in (D); n = 5; error bars represent S.E.M. (H) RNaseH^D10R-E48R^-mCherry accumulation at laser damage sites was measured in wild-type, CtIP-depleted, XPG-depleted, or both CtIP/XPG-depleted U2OS cells as in (D); n > 8, error bars represent S.E.M.

Based on the yeast experiments with *SAE2*, we hypothesized that overexpression of a helicase that is specific to RNA-DNA hybrids should rescue the sensitivity of CtIP-depleted cells to CPT exposure. To test this idea, we complemented the CtIP-depleted cells with the C-terminal domains (a.a. 1851-2677) of human Senataxin, the ortholog of yeast SEN1 that also acts to remove RNA-DNA hybrids and promotes transcription termination ^37^. Remarkably, overexpression of Senataxin in the CtIP-depleted cells completely rescues the CPT sensitivity; however, unlike the yeast experiments, no effect of PCF11 expression was detected (Fig. 4A). Although Senataxin expression rescued the survival of CtIP-depleted cells after CPT exposure, only partial suppression of the defects in growth and DNA end resection were observed (Fig. S2C, D). These results also suggest that inhibition of transcription could rescue the CPT sensitivity of CtIP-depleted cells. Indeed, we observe that pre-treatment of the cells with DRB, an inhibitor of RNA polymerase II-dependent RNA synthesis, partially rescues the CPT sensitivity caused by CtIP deficiency (Fig. 4B). Thus, similar to yeast, reduction of transcription alleviates the effects of CtIP deficiency in human cells, suggesting a conserved mechanism for the repair of Top1 lesions associated with aberrant transcription.

### CtIP deficiency induces accumulation of R-loops at laser micro-irradiation sites

Similar to topoisomerase-DNA adducts, UVA laser-induced DNA crosslinks present a physical barrier to RNA polymerase that could stall transcription and promote the formation of complex DNA lesions including R-loops. We utilized a live cell assay with lentiviral expression of a bacterial RNaseH catalytic mutant fused to mCherry (RNaseH^D10R-E48R^-mCherry) that acts as a sensor for R-loops in the genome ^58^. The RNaseH^D10R-E48R^-mCherry sensor was expressed in the wild-type or CtIP-depleted U2OS cells described above, which were laser micro-irradiated in a small area of the nucleus. Measurement of the accumulation of RNaseH^D10R-E48R^-mCherry signal over time in these cells showed that the recruitment of RNaseH occurs to a higher intensity in CtIP-depleted cells in comparison to cells complemented with wild-type CtIP, suggesting that higher levels of R-loops are formed in the absence of CtIP (Fig. 4C, D). A similar control experiment examining mCherry alone did not show this pattern, confirming the effect is specific to the RNaseH fusion (Fig. S3). Since we found that overexpression of Senataxin rescues the CPT sensitivity of CtIP-depleted cells, we reasoned that it might also rescue the higher levels of R-loop accumulation in the absence of CtIP. We expressed the C-terminal domains of human Senataxin in the CtIP-depleted cells, and measured the levels of laser-induced R-loops by the live cell imaging method. Here also we found that with Senataxin expression, the recruitment of RNaseH in the CtIP-depleted cells is reduced to the levels observed in cells expressing wild-type CtIP (Fig. 4D). Similar to the CPT survival assay results, we found the PCF11 was not able to rescue CtIP deficiency for reduction of R-loop levels after laser micro-irradiation.

Previous work on CtIP in vitro identified mutants that exhibit lower levels of endonuclease activity on flap structures in comparison to the wild-type protein ^14,15^. Here we used the N289A/H290A (NA/HA) mutant allele also containing shRNA-resistant mutations to complement CtIP-depleted cells and found that this mutant failed to reduce the increased RNaseH recruitment to damage sites that occurs in CtIP-deficient cells (Fig. 4E, S2E). Thus the nuclease activity of CtIP appears to play a role in DNA damage recognition and/or processing that helps to prevent R-loop accumulation in human cells.

### The accumulation of R-loops in CtIP deficient cells is dependent on transcription

To further test the role of active transcription in R-loop accumulation, we pre-treated cells with 5,6-dichloro-1-beta-D-ribofuranosylbenzimidazole (DRB), a RNA polymerase II inhibitor, and found that inhibition of transcription rescues the R-loop accumulation phenotype caused by CtIP deficiency (Fig. 4F). As overexpression of Senataxin reduces R-loop formation in CtIP-depleted cells after DNA damage, we expected that depletion of this enzyme would have the opposite effect. Indeed, U2OS cells depleted of Senataxin also exhibit increased R-loop formation after laser-induced DNA damage (Fig. 4G, S2F), consistent with previous observations of DNA damage sensitivity in Senataxin mutant cells ^59,60^.

The XPG protein, a component of nucleotide excision repair ^61^, has been shown to be important in the resolution of R-loops ^62^. As a comparison to CtIP, we also assessed whether the deficiency of this protein would also lead to high R-loop levels after DNA damage. As expected, we observed that the XPG deficient cells accumulate more R-loops after laser induced DNA breaks in comparison to wild-type cells (Fig. 4G, S2G). Interestingly, however, concurrent depletion of both CtIP and XPG showed R-loops at levels comparable to wild-type untreated cells (Fig. 4H). This result suggests that R-loops do not form efficiently in the absence of both CtIP and XPG, or that they are not recognized efficiently by the RNaseH^D10R-E48R^-mCherry protein in the absence of both CtIP and XPG.

### R-loop accumulation in CtIP-depleted cells without exogenous damage

Considering that deletion of the gene encoding CtIP is cell-lethal even in the absence of exogenous damage ^63,64^, we also hypothesized that R-loops might accumulate in CtIP-depleted cells under normal growth conditions. To address this question, we again used the RNaseH^D10R-E48R^-mCherry sensor but in this case monitored its accumulation in undamaged cells by fluorescence activated cell sorting (FACS), using a modified technique reported previously with a fragment of RNaseH fused to GFP ^58^. Using this procedure, unbound RNaseH^D10R-E48R^-mCherry protein is removed from the nucleoplasm by detergent extraction, while protein bound to chromatin is retained. This analysis showed a statistically significant increase in RNaseH^D10R-E48R^-mCherry fluorescence intensity in CtIP-depleted U2OS cells in all cell cycle phases (Fig. 5A). Analysis of mCherry alone in these cells showed no differences with CtIP depletion (Fig. S3). This was also observed in XPG-depleted cells, consistent with the observed effects of both XPG and CtIP with laser damage as shown in Fig. 4. Similarly, we examined R-loop accumulation in CtIP-depleted cells complemented with the nuclease-deficient form of CtIP (N289A/H290A, NA/HA, Fig. S3D) and found the levels of RNaseH^D10R-E48R^-mCherry fluorescence intensity identical to that of CtIP-depleted cells (Fig. 5B), suggesting that the nuclease activity of CtIP is necessary for R-loop resolution even in cells that are not exposed to exogenous damage.

**Figure 5.**
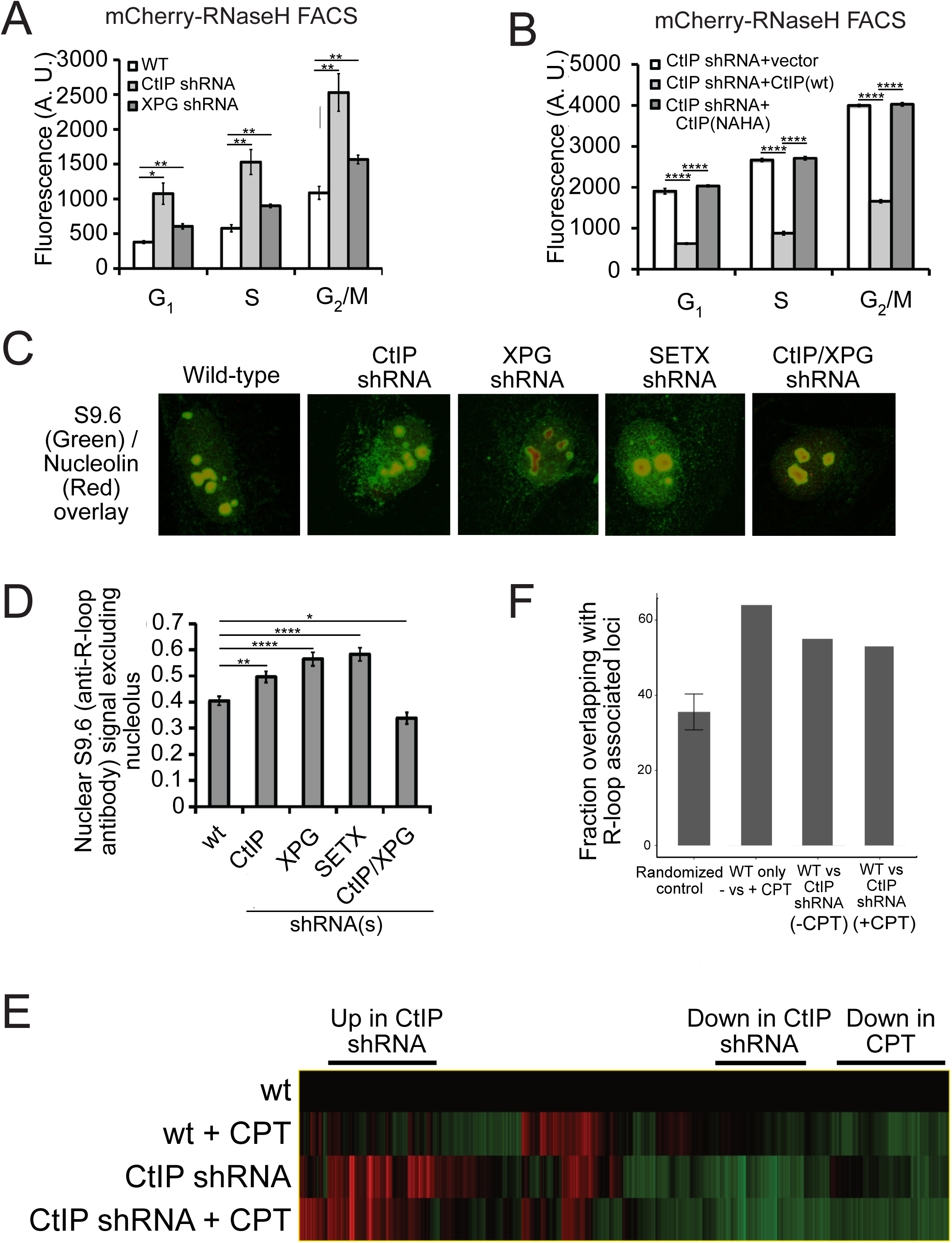
CtIP depletion affects R-loop accumulation and transcription in human cells. (A) DNA-RNA hybrids were quantified in undamaged U2OS cells by monitoring chromatin-bound RNaseH^D10R-E48R^-mCherry by FACS. 10,000 cells were counted in each of 3 biological replicates; error bars represent S.D. * and ** denote p<0.05 or 0.01, respectively in Student’s two-tailed T test with comparisons as indicated. (B) RNA/DNA hybrids were quantified in undamaged, CtIP-depleted U2OS cells complemented with either vector only, eGFP-CtIP(wt), or nuclease-deficient eGFP-CtIP(NA/HA) as in (A). 10,000 cells were counted in each of 3 biological replicates; error bars represent S. D. **** denotes p< 0.0001 using Student’s two-tailed T test with comparisons as indicated. (C) S9.6 antibody was used to monitor RNA-DNA hybrids in wild-type or CtIP-depleted U2OS cells. Anti-Nucleolin was used as a marker for the nucleolus. (D) Quantification of S9.6 antibody signal in wild-type, CtIP-depleted, XPG-depleted, Senataxin-depleted, or double CtIP/XPG-depleted U2OS cells as indicated. Signal overlapping the nucleolin signal was excluded from the analysis; n > 50, Error bars represent S.E.M. *, **, and **** denote p< 0.05, 0.01, and 0.0001, respectively, using Student’s two-tailed T test with comparisons as indicated. (E) Wild-type or CtIP-depleted U2OS cells were exposed to CPT (5 µM) or were untreated before harvesting of cellular mRNA. Analysis of transcripts by RNA-seq and hierarchical clustering of transcripts from 21,412 genes is shown as a heat map (red for over-expressed, black for unchanged expression, and green for under-expressed genes) in comparison to wild-type undamaged cells (see Table S3). (F) Statistical comparisons of overlap between the top 100 differentially expressed genes as ranked by DESeq differential expression p-value and DRIPc-seq peaks from GEO dataset GSE70189. Randomly picked genes were compared to this dataset (“randomized control”) with the average of 35.56% and standard deviation 4.76% (estimated from 1000 simulations). Genes with significant differences between wild-type and CtIP-depleted cells were identified in the absence of DNA damage (“WT vs CtIP shRNA (-CPT”)) as well as with CPT treatment (“WT vs CtIP shRNA (+ CPT”)) and were compared with the DRIPc-seq dataset. All 3 values are above the 99% confidence intervals for the null hypothesis that the genes showing the most evidence of differential expression overlapped the peak regions at the same rate as randomly selected genes.

To confirm these results with a different technique, we also utilized the S9.6 antibody, which recognizes RNA-DNA hybrids and has been widely used as a probe for R-loops in cells ^26,53,65^. We used immunofluorescence signal from S9.6 antibody in control as well as CtIP-depleted, XPG-depleted, and Senataxin-depleted U2OS cells and quantified the level of signal per cell. As this antibody also recognizes the components of nucleoli, the S9.6 signal that colocalized with these organelles was subtracted from the total signal, and the result was normalized by the area of the nucleus. We found that undamaged CtIP-depleted or XPG-depleted cells have significantly more R-loops than their wild-type counterparts (Fig. 5C, D). These findings suggest that CtIP is responsible for the prevention and/or resolution of R-loops, in normally growing cells as well as in cells exposed to DNA damage. We also observed that concurrent depletion of CtIP and XPG resulted in a lower level of R-loop accumulation (Fig. 5D), similar to the result with laser-induced damage (Fig. 4H). This observation of lower RNA-DNA hybrids in the absence of both nucleases is thus not specific to laser damage or to the sensor used for R-loop detection.

### Global patterns of transcription are altered with CtIP depletion

CtIP was originally identified as a binding partner of C-terminal Binding Protein (CtBP), a transcriptional co-regulator **^66^**, and also binds directly to Rb and the tumor suppressor BRCA1 which also has been widely reported to affect transcription and R-loop formation ^67–71^. Considering our results with CtIP and R-loop accumulation, we asked whether depletion of CtIP has global effects on transcription patterns by analyzing mRNA using RNA-seq. We performed this analysis on U2OS cells expressing control or CtIP-specific shRNA and examined both CPT-treated and untreated conditions. RNA levels were quantified for 30,769 transcripts. CtIP depletion was found to alter 5,013 (∼ 16%) of these transcripts, with both increases (2,578) and decreases (2,435) observed relative to the control cells. Unsupervised hierarchical clustering was used to analyze the transcripts (Fig. 5E) (Table S3). We found that there are transcriptional changes associated with CPT exposure in U2OS cells: 1,285 transcripts were upregulated upon CPT treatment, and 1,158 transcripts were downregulated. Interestingly, a comparison of the genes affected by CPT exposure to a previous dataset of genes showing R-loop accumulation indicates an overlap significantly higher than would be predicted by chance (Fig. 5F). Over 60% of the genes affected by CPT (either up or down) overlap with R-loop prone regions of the genome **^72^** (see “WT only, - vs + CPT”, Fig. 5F), whereas less than 40% overlap with DRIP positive genes occurs with a randomized control. A similar analysis of the gene set identified in CtIP-depleted cells compared to normal cells also indicates a higher than expected overlap (∼50%), suggesting that depletion of CtIP has effects on transcription that are correlated with R-loop-prone regions of the genome.

### Quantitation of RNA-DNA hybrids in CtIP-depleted cells

To investigate the role of CtIP in R-loop accumulation at specific loci, we first generated an inducible genomic cassette containing a mouse class switch region that has previously been shown to accumulate R-loops ^73^. We performed DNA-RNA immunoprecipitation (DRIP) with the S9.6 antibody followed by quantitative PCR for this locus and found that the levels of R-loops increase with doxycycline-induced transcription, even in wild-type cells (Fig. 6A). The DRIP signal was removed by RNase H treatment, confirming the specificity of the S9.6 antibody. A much larger increase in R-loops was found in CtIP-depleted cells, however, while the overall levels of transcripts are similar in both cases (Fig. 6A, Fig. S4A).

**Figure 6.**
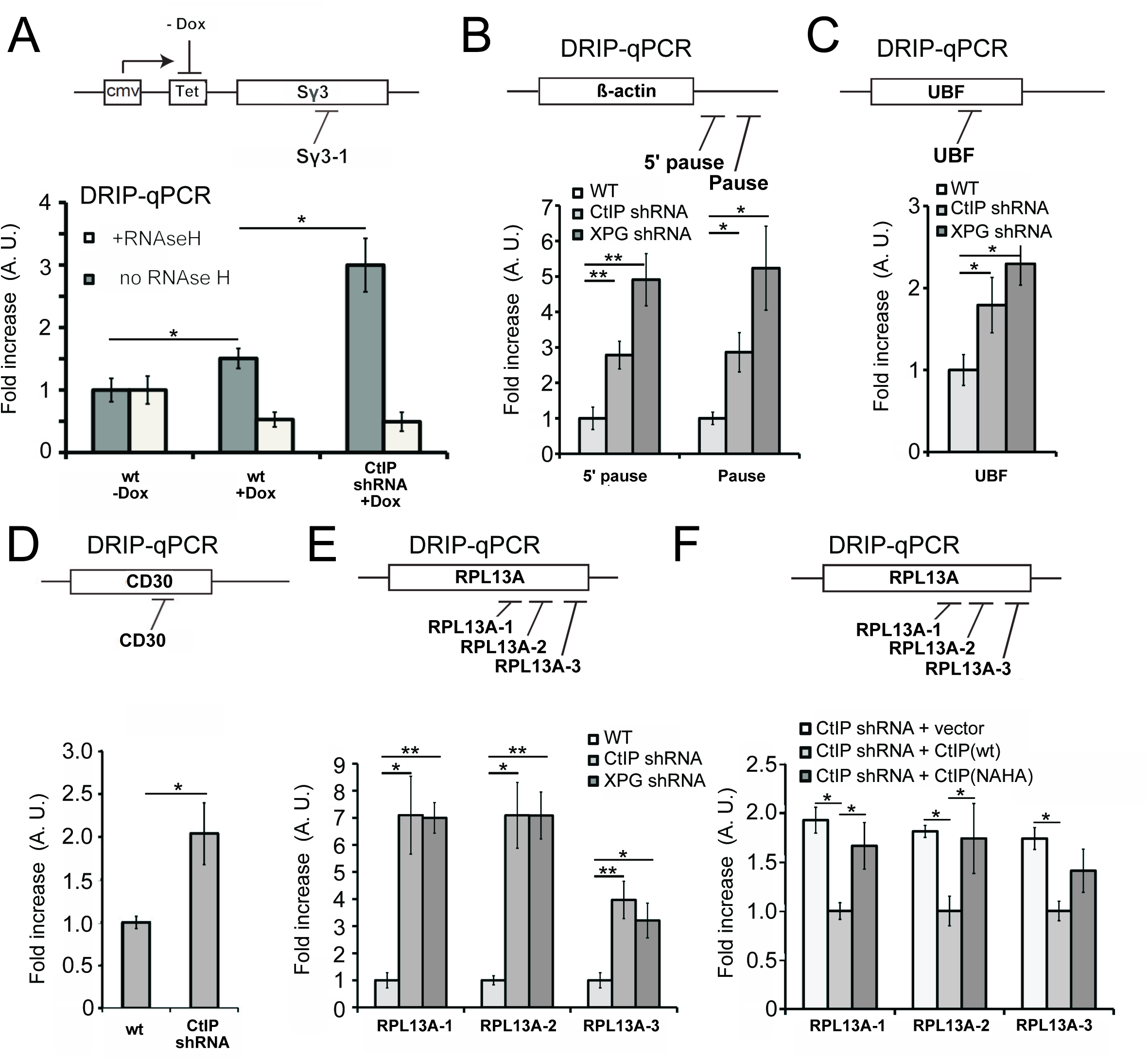
CtIP depletion induces accumulation of R-loops in human cells. (A) RNA/DNA hybrids were quantified by DRIP-qPCR in U2OS cells containing a stably integrated, doxycycline-inducible transgene containing murine Sγ3 repeats. R-loop accumulation was measured by immunoprecipitation with the S9.6 antibody and qPCR for the Sγ3 region in the absence or presence of transcription (-/+ Dox) in wild-type or CtIP-depleted cells; n > 6, error bars represent S.E.M. (B), (C), (D), (E), (F) Quantification of RNA/DNA hybrids in U2OS cells at endogenous loci using DRIP-qPCR. Levels of hybrids were measured at the ß-actin (B), UBF (C), CD30 (D), RPL13A (E, F) genes; n > 3, error bars represent S. D. In (F), CtIP-depleted cells were complemented with wild-type eGFP-CtIP or nuclease-deficient NA/HA mutant. * and ** denote p< 0.05 or 0.01, respectively, using Student’s two-tailed T test with comparisons as indicated.

We then examined several endogenous loci that have previously been reported to be prone to R-loop formation ^67,74^. The β-actin gene has been reported to accumulate R-loops, particularly at the G-rich “pause” sequences downstream of the coding region ^67,75^. We observed higher levels of RNA-DNA hybrids at these regions in both CtIP-depleted as well as XPG-depleted cells during normal cell growth (Fig. 6B). Similar results were observed on the UBF and CD30 genes (Fig. 6C, D), which we examined because levels of transcription are significantly lower in CtIP-depleted cells (Table S3, Fig. S4). We also examined RPL13A, a gene that has been identified as a region prone to R-loop formation ^65,74^. R-loop formation was 3 to 7-fold higher in CtIP-depleted as well as in XPG-depleted cells compared to wild-type, measured at 3 locations throughout the body of the gene (Fig. 6E). The nuclease-deficient NA/HA allele of CtIP was expressed in CtIP-depleted cells, and R-loops were found to be approximately 1.5 to 2-fold higher in these cells in comparison to cells expressing the wild-type allele, similar to our observations in uncomplemented cells (Fig. 6F).

### CtIP depletion leads to fewer DNA breaks after CPT treatment

Our results suggest that the CtIP and XPG nucleases help to either prevent R-loop formation in the genome or to resolve R-loops once they are formed. Since CtIP and XPG are both specific for 5’ flaps, we considered a model in which CtIP and XPG process 5’ flaps present in an R-loop structure (Fig. 7). We hypothesize that this ssDNA cleavage event would result in extension and stabilization of the RNA-DNA hybrid, because the release of tension in one DNA strand would prevent spontaneous extrusion of the RNA (Fig. S5). This conversion of the nascent lesion into an extended structure would also generate access for helicases like Senataxin to recognize and remove the RNA strand from the DNA. It is also possible that nuclease processing could convert the pre-lesion into double-strand breaks.

**Figure 7.**
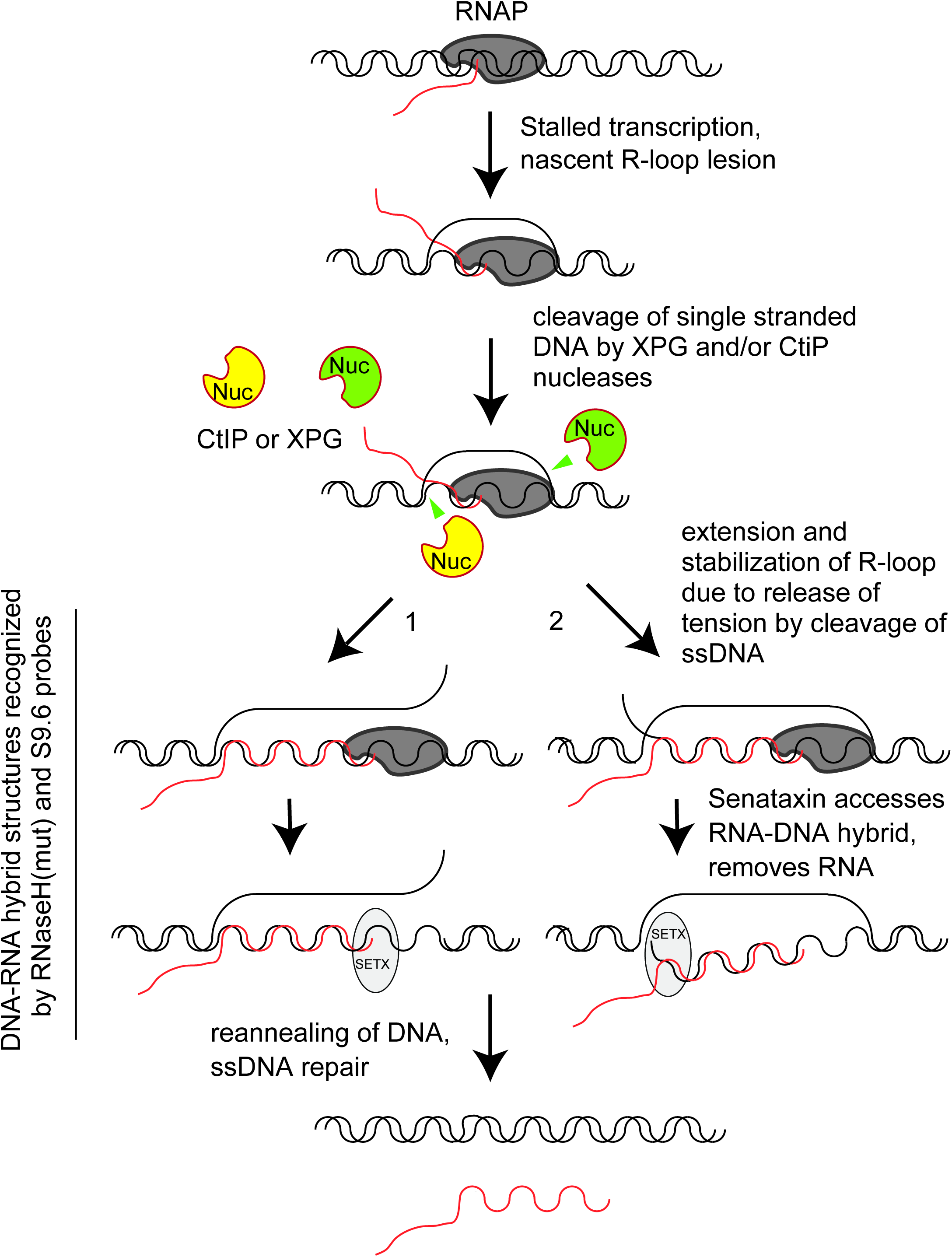
A model of R-loop processing. Polymerase stalling at sites of nicks, topoisomerase adducts, or other lesions generates a nascent R-loop structure. Cleavage of this nascent structure at one of 2 exposed 5’ flaps by CtIP or XPG (cutting of non-template or template strand; 1 or 2, respectively) can generate a stabilized, extended RNA-DNA hybrid because of the release of torsional constraint in the DNA template. Senataxin (or other helicases) preferentially access the RNA-DNA hybrid in this stabilized intermediate and remove the RNA, promoting reannealing of the displaced non-template strand and single-strand DNA repair.

One prediction of this model is that CtIP and XPG-depleted cells would exhibit fewer single-strand DNA breaks compared to normal cells, even though they exhibit higher levels of R-loops. To test this idea, we created DNA damage with CPT and measured single-strand DNA breaks with an alkaline comet assay. We found that exposure to CPT generated a marked increase in DNA breaks, and that transcription promotes these breaks, as pretreatment with DRB reduced the levels of breaks by ∼75% in wild-type cells (Fig. 8A). CtIP depletion led to a significant reduction in the accumulation of DNA breaks after CPT treatment, consistent with the proposed model that CtIP can facilitate the conversion of stalled transcription associated DNA lesions into DNA breaks.

**Figure 8.**
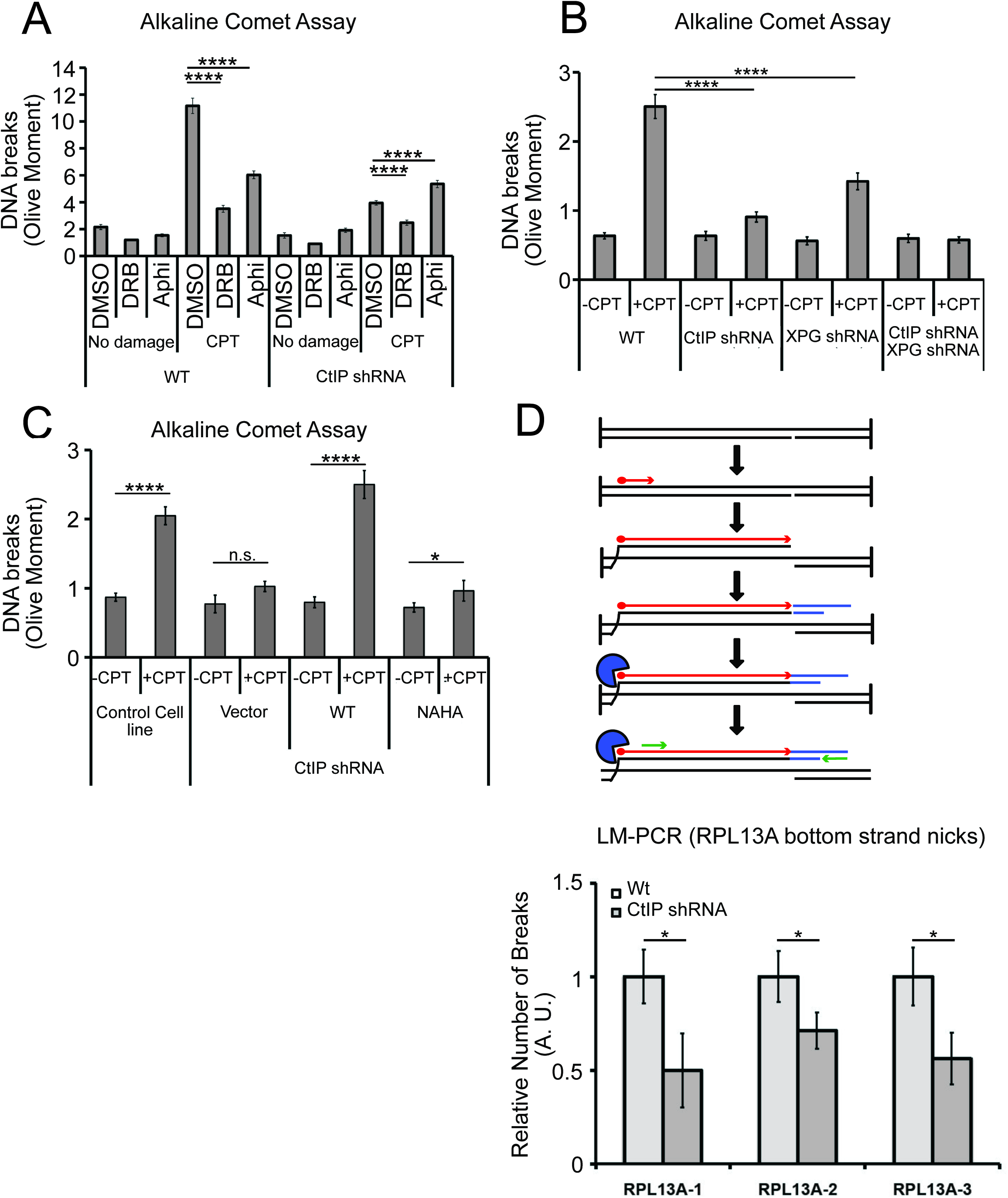
CtIP and its nuclease activity promote ssDNA break formation. (A) DNA breaks were quantified in wild-type and CtIP-depleted U2OS cells by alkaline comet assay. Cells were untreated (DMSO) or exposed to 5 µM CPT for 1 hr, with DRB (20 µM) or aphidicolin (2 µg/mL) pretreatment as indicated. Olive moments were calculated by analyzing at least 100 comets for each sample; error bars represent S.E.M. (B) Quantification of ssDNA breaks by alkaline comet assay was performed in wild-type, CtIP-depleted, XPG-depleted, or CtIP/XPG-depleted U2OS cells as in (A). (C) Quantification of ssDNA breaks by alkaline comet assay was performed in wild-type or CtIP-depleted U2OS cells complemented with wild-type eGFP-CtIP or nuclease-deficient NA/HA mutant CtIP as in (A). (D) A schematic representation of Ligation-mediated (LM)- PCR assay (top). Primer extension with a biotinylated primer (in red) from genomic DNA produces a double-stranded DNA end that is isolated with streptavidin and amplified by ligation-mediated PCR (asymmetric duplex and nested primers are presented in blue and green, respectively). LM-PCR assay measuring DNA breaks on the bottom strand of the RPL13A gene (bottom). DNA single-strand breaks were measured by LM-PCR at the endogenous RPL13A locus in wild-type or CtIP-depleted cells; n = 6, error bars represent S.E.M. * and **** denote p< 0.05 or 0.0001, respectively, using Student’s two-tailed T test with comparisons as indicated.

Next, we assessed the levels of single-strand DNA breaks in cells depleted for XPG, or for XPG and CtIP concurrently. In cells depleted for XPG there are fewer breaks after CPT treatment, and in cells depleted for both proteins, we observed no increase in breaks at all with CPT exposure (Fig. 8B). This result does suggest that XPG and CtIP perform a function at transcription-associated lesions that results in an increase in single-strand breaks, likely their common function in promoting cleavage of 5’ flaps. To test this, we also examined breaks by comet assay in cells expressing the nuclease-deficient NA/HA CtIP mutant and found that in this case also, there was a large reduction in breaks formed with CPT exposure, very similar to the uncomplemented cells (Fig. 8C).

To further validate this result at a specific locus in the genome, we returned to the RPL13A gene, where we observed high levels of R-loops in cells lacking CtIP using DRIP-qPCR (Fig. 6). Here we used a modified LM-PCR assay ^67^ to measure single-strand DNA breaks in the genome. Primer extension from a sequence within the RPL13A locus converts any single-strand breaks present into single-ended double-strand breaks, then ligation-mediated PCR followed by qPCR quantitates the levels of these single-ended breaks (Fig. 8D). Using this procedure, we did observe spontaneous single-strand breaks on the template strand of the RPL13A gene, and the level of breaks was reduced by 40 to 50% with depletion of CtIP, again consistent with the observation that single-strand breaks are reduced in cells lacking CtIP.

## Discussion

In this study we demonstrate a functional relationship between the Sae2/CtIP enzyme and the RNA-DNA helicase SEN1/Senataxin in eukaryotic cells, showing that cells lacking either enzyme are hypersensitive to DNA damage, and that overexpression of SEN1/Senataxin can rescue the damage sensitivity of cells lacking Sae2/CtIP. We further provide evidence for accumulation of stalled RNA polymerase complexes and RNA-DNA hybrids in cells deficient in Sae2/CtIP, with these lesions located at sites of highly expressed genes. These results suggest that Sae2/CtIP is not only a double-strand break resection factor, but is a sensor and processing enzyme for transcription-associated lesions in eukaryotic cells.

In *S. cerevisiae*, the temperature-sensitive *sen1-1* mutant exhibits synthetic lethality in combination with DNA repair mutants *mre11, rad50, sgs1*, and *rad52*, indicating a requirement for homologous recombination in the absence of *SEN1* function ^30^. Consistent with these results, we observe a synthetic sensitivity of *sen1-1* with *sae2* for survival of CPT and MMS exposure. Conversely, overexpression of *SEN1*, and to a lesser extent *PCF11*, rescues *sae2* survival of these DNA damaging agents.

From *sen1* separation of function mutants we conclude that the helicase activity of *SEN1* is important for the effect on survival of *sae2* strains to DNA damage. Since we also observe stalling of RNA polymerase in *sae2* strains, particularly with CPT treatment, the simplest explanation is that Sae2 promotes the processing or resolution of transcription complexes stalled by Top1 conjugates and that *SEN1* overexpression rescues survival in this context because of its known activities in removing RNA-DNA hybrids.

Although replication definitely contributes to CPT toxicity in this context, we found that the sites of Sae2 accumulation during DNA damage exposure are highly expressed genes, not replication origins. This observation coincides with the known propensity for Top1 to accumulate at sites of active transcription and to facilitate transcription elongation ^76–78^.

In human cells, overexpression of Senataxin also complements CtIP-depleted cells for CPT survival and for accumulation of R-loops at sites of laser-induced DNA damage. The extent of this suppression is remarkable and suggests that R-loops are a limiting factor in the recovery of CtIP-depleted cells following DNA damage. In mammalian cells, loss of CtIP is lethal whereas yeast strains lacking *SAE2* have no growth deficit in the absence of DNA damage ^22,24,55,63^. This could be related to our observation that CtIP-depleted human cells exhibit higher levels of spontaneous R-loops than wild-type cells, even in the absence of exogenous damage, whereas *sae2* null yeast strains only show RNA polymerase II pausing and R-loop accumulation with damage treatment.

CtIP and Sae2 exhibit an intrinsic endonuclease activity that is specific for 5’ flap structures and can be genetically separated from its ability to stimulate the nuclease of activity of Mre11 ^79^. In human cells we observed that a CtIP mutant deficient in endonuclease activity failed to rescue CtIP-depleted cells for reduction of R-loops at laser damage sites or at genomic loci, thus we conclude that the action of CtIP at sites of stalled transcription involves its nuclease activity. Since we also show in this work that R-loop formation in CtIP-depleted cells resembles that of cells depleted of XPG, an endonuclease that also acts in nucleotide excision repair, the simplest hypothesis is that both enzymes target 5’ flap structures, of which there are at least two present in every R-loop structure (see model in Fig. 7). Evidence for a DNA cleavage event induced by CtIP or XPG in response to CPT or other transcription stalling lesions comes from our analysis of CPT-treated cells using alkaline comet assays, and also by locus-specific quantitation of single-strand breaks. These results clearly show that there are fewer single-strand breaks after DNA damage in the absence of CtIP or XPG, even though the levels of R-loops in these cells are much higher. We do not know if there is a specificity for either template or non-template ssDNA breaks for these enzymes, but we do observe a reduction in template strand breaks at the RPL13A locus with depletion of CtIP.

Although R-loops as measured by RNaseH-binding and S9.6 antibody immunoprecipitation are significantly higher in CtIP or XPG-depleted cells, depletion of both factors simultaneously shows a very different result: R-loop levels similar to wild-type cells. We observed this phenomenon in untreated cells, in cells treated with CPT, and in cells with laser-induced DNA damage. To explain this finding, we propose that CtIP and XPG are functionally redundant in their ability to process R-loop structures; thus in their absence, unprocessed nascent R-loops accumulate (Fig. 7). We propose that, due to the topological constraints on R-loop formation, these nascent R-loop structures are small and are not efficiently recognized by either the S9.6 antibody or by RNase H. We know that there are toxic lesions present in cells depleted of both CtIP and XPG since the survival of cells depleted for both factors is very poor (Fig. S2H).

This hypothesis is attractive from a topological standpoint, because a nascent R-loop is extremely constrained. Lieber and colleagues have defined the area of nascent R-loop formation as the “R-loop initiation zone”, where clusters of G residues in the nontemplate strand promote the start of R-loop hybridization and increase the likelihood that extensive R-loop formation occurs in the “R-loop elongation zone” ^80^. This study also showed that negative supercoiling can reduce the necessity for G clusters in the initiation zone, and a nick in the non-template strand greatly improves the efficiency of R-loop formation, presumably because the non-template strand then does not compete with the RNA for hybridization to the template strand. Here we postulate that transient stabilization of RNA by annealing with the template DNA strand could initiate R-loop formation but that extensive spreading and stabilization of the RNA-DNA hybrid would require either secondary structure formation in the non-template strand or cleavage of one of the DNA strands as shown by Roy et al. We propose that cleavage of one of the DNA strands, although seemingly detrimental in its stabilization of the RNA-DNA hybrid, likely facilitates removal of the hybrid by transcription-associated helicases such as Senataxin or Aquarius. Biochemical characterization of the helicase domain of yeast Sen1 showed that the enzyme requires single-stranded DNA adjacent to the hybrid for efficient loading prior to unwinding of the annealed RNA ^81^. This type of structure is not available in an R-loop in a topologically closed system without cleavage of a DNA strand or active unwinding of the DNA duplex adjacent to the RNA-DNA hybrid. In addition, a recent study found that treatment of chromatin with low levels of the single-strand DNA-specific S1 nuclease greatly improves the yield of R-loops immunoprecipitated by the S9.6 antibody ^82^, consistent with this hypothesis.

In conclusion, we have presented evidence supporting a role for Sae2/CtIP in processing of transcription-related DNA lesions and show that release of R-loops from the genome is a limiting factor for the survival of Sae2/CtIP-deficient cells following DNA damage. This is perhaps not so surprising as CtIP was first identified through its association with the transcription regulator CtBP ^66^, and also interacts directly with the tumor suppressor BRCA1, which also associates with transcription-related complexes ^55,71,83^. BRCA1 itself has been shown to localize to a subset of transcription termination regions and to regulate levels of RNA-DNA hybrids in human cells ^67,84^. In addition, XPG was recently found to be present in a complex with BRCA1 that is important for cellular responses to DNA damage that is separate from its roles in excision repair ^85^. Further studies will need to address whether the roles of CtIP and BRCA1 at transcription lesions are functionally interdependent and whether Senataxin can also rescue BRCA1-deficient cells in the same manner as CtIP-depleted cells. Lastly, CtIP has been shown to have important repair functions in G_1_ phase cells ^86–90^; it will be important to determine if the R-loop processing functions indicated here play a role in the repair of transcription-associated damage even outside of S phase and whether this accounts for any of the essential functions of CtIP in mammals.

## Materials and Methods

### S. cerevisiae strains and expression constructs

For yeast strains see Supplementary Table 1. Genomic deletions were made with standard lithium chloride transformations and integrations ^91^ and were verified by PCR. Full-length *SEN1, SSU72*, and *RTT103* were cloned into the 2μ plasmid pRS425 ^92^ by fusion PCR to create pTP3500, pTP3249, and pTP3498, respectively. A pRS425 derivative containing *PCF11* was a gift from Steve Hanes. Mutations in *SEN1* were made in pTP3500 by Quikchange mutagenesis (Stratagene). The *sen1-1* strain and corresponding wild-type strain were gifts from Nicholas Proudfoot. A high-copy vector containing the wild-type *SAE2* gene with a 2xFLAG tag in pRS425 (61)“*FLAG-SAE2*/2μ” (28) was a gift from John Petrini, and was used for the *SAE2* ChIP experiment.

### Yeast Camptothecin Survival Assay

Cells were grown to exponential phase (OD 0.5-0.6) and synchronized using alpha-factor for 3 hrs at 30°C. Half of the cells were kept in alpha factor (G_1_), while the other half were washed and resuspended in fresh media (S). Previous studies have shown that all the cells enter S phase 30 min after the removal of alpha factor ^93^. Therefore 30 min after the release into fresh media, both G_1_ and S samples were treated with either 100 µM CPT or DMSO for two hours for an acute drug dose. At the end of the treatment, cells were washed and appropriate dilutions were plated on YPDA + glucose plates to obtain single colonies. Single colonies were counted after two days and survival efficiency was calculated as the number of colonies on the “+ CPT” plates divided by the number of colonies on the “no treatment” plates. For thiolutin treatment, 15 min after the release into fresh media both G_1_ and S phase cells were treated with either DMSO or thiolutin (2.5 μg/mL final concentration). After another 15 min CPT or DMSO was added to the media at indicated concentration.

### Chromatin Immunoprecipitation in yeast

Sae2: Yeast cells containing a FLAG-Sae2 expression cassette on a high copy number 2μ plasmid were grown to exponential phase in 2 L of appropriate minimal media to OD_600_ = 0.5-0.6. The cells were centrifuged and resuspended in 50ml fresh media (concentrated 20-fold). Yeast mating pheromone alpha-factor was added to final concentration of 10mM and cells were synchronized for 4 hrs at 30°C. Half of the cells were kept in alpha factor (G_1_), while the other half were washed and resuspended in fresh media (S). After the wash, both G_1_ and S phase cells were further divided into two and treated with 100 μM camptothecin or DMSO for 50 min. At the end of the drug treatment cells were crosslinked by addition of formaldehyde (1% final concentration) at RT for 25 min, followed by glycine (125 mM final concentration) at RT for 5 - 10min. The cells were harvested, washed with water and flash frozen until further processing. Thus 2L of starting culture was divided into 4 samples per strain with approximately 500 OD cells per sample.

Each cell pellet was resuspended in 400 μl lysis buffer (50 mM HEPES, 150 mM NaCl, 1 mM EDTA, 1 mM DTT, 10% glycerol, 0.1% NP40, and protease inhibitors (Pierce # A32955; 1 tablet per 10 ml) and lysed using a bead beater (3 cycles of 45 sec) in the presence of 400 μl 0.5 mm zirconia beads. The beads were washed with 2 x 500μl of lysis buffer to collect a total of 1.5ml cell extract. Sonication was performed with a Branson Digital Sonifier (5 cycles of 1 min each, with 10 sec on, 10 sec off at 28% amplitude). The cells were kept on ice for 5 min in between each cycle. The extract was cleared with high-speed centrifugation at 4°C for 30 min. A small sample was treated with proteinase K and RNaseA and separated by agarose gel to check for sonication efficiency (DNA size should be between 100-250 bp). The lysate was diluted to 10 ml with the lysis buffer. A 500 μl aliquot of the lysate was saved as the “Input” sample. 50 μg Sigma FLAG-M2 antibody and 300 μl Pierce Protein A/G magnetic beads (prewashed with lysis buffer) were added to the cleared lysate. The cells were rotated at 4°C overnight. The following day, the beads were separated and washed 3 times each with wash-buffer 1 (25 mM Tris pH 8, 1 mM EDTA, 150 mM NaCl, 1% Triton, 100 μM camptothecin or DMSO equivalent), wash buffer 2 (25 mM Tris pH = 8, 1 mM EDTA, 500 mM NaCl, 1% Triton), wash buffer 3 (10 mM Tris pH 8, 1 mM EDTA, 1% Triton, 500 mM LiCl, 0.5% NP40) and wash buffer 4 (25 mM Tris pH 8, 1 mM EDTA, 1% Triton, 150 mM NaCl). 500μl elution buffer (wash buffer 4 + 3X FLAG peptide) was added to the beads and kept shaking at 4°C for 2h. The beads were eluted with 100 μl elution buffer (wash buffer 4 + 2% SDS) and incubation at 65°C for 30 min. A second elution was done with 100 μl elution buffer similarly and combined for 150 μl total elution. The final elution and the Input sample (supplemented to 2% final SDS) were reverse cross-linked at 65°C overnight. Next day, all samples were diluted to SDS < 0.5% and treated with 20 μg RNase A at 37°C 2h followed by 40 μg of proteinase K at 37°C 2h. The DNA was purified by phenol chloroform extractions followed by ethanol precipitation and stored at - 20°C until further processing.

The DNA samples were gel purified to obtain a size range of 50-200bp. The sequencing libraries were prepared for each sample using NEBNext DNA Library prep kit for Illumina (E6040S). and were sequenced and analyzed at the New York Genome Center. The bam files and bed files were visualized using Integrated Genome Viewer (iGV) and compared with saccer3 reference genome.

Rpb2: Yeast cells containing a His6-Tev-Biotin (HTB)-tagged Rpb2 allele in the genome were used for this experiment. Cell pellets were prepared, lysed and sonicated similar to Sae2 ChIP, except the following lysis buffer was used - 25 mM Tris pH 7.5, 150 mM NaCl, 1 mM EDTA, 1 mM DTT, 10% glycerol, 0.1% NP40, 0.1% SDS, 1% Triton, 200 mM PMSF supplemented with protease inhibitors (Pierce # A32955; 1 tablet per 10 ml). After sonication the lysate was divided into three parts - 10% for Input, 45% for Rpb2 IP and 45% for mock IP. The IP was done with streptavidin linked M280 magnetic beads (Invitrogen) overnight and sheep anti-mouse M280 magnetic beads (Invitrogen) were used for the mock. The next day the beads were separated and washed 3 times with the following buffers - 1 (25mM Tris pH 7.5, 150 mM NaCl, 1 mM EDTA, 1% Triton, 0.1% SDS), 2 (25 mM Tris pH 7.5, 500 mM NaCl, 1 mM EDTA, 1% Triton, 0.1% SDS), 3 (10 mM Tris pH 8, 500 mM LiCl, 1mM EDTA, 1% Triton, 0.5% NP40), and 4 (10mM Tris pH 8, 1mM EDTA). For each wash, the beads were rotated for 5 min at RT. The fold enrichment over Input at different genomic loci was calculated using SYBR-Green based quantitative PCR. Samples were tested at various dilution factors to optimize a dilution factor for a CT value in the linear range. Then percentages of DNA enrichment of IP over the Input were calculated by using the following equation:

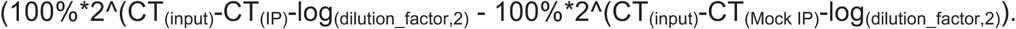

Then relative fold of enrichment was calculated by normalizing the percentage of DNA enrichment of IP of each group to WT cells without CPT treatment.

### DRIP-qPCR in yeast

Yeast cells were grown as described for the Sae2 ChIP experiment, except there were no G2 cell samples and no formaldehyde crosslinking was done. DRIP was performed according a protocol from Dr. Douglas Koshland ^82^. A detailed protocol is available on request. Briefly, 150–200 μg genomic DNA isolated using the Qiagen Genomic DNA kit (500G) was treated with S1 nuclease and precipitated in 130μl TE. The DNA was sonicated using a Covaris sonicator, precipitated, and resuspended in 50 μl of nuclease-free water. Then 350μl of FA buffer (1% Triton X-100, 0.1% sodium deoxycholate, 0.1% SDS, 50 mM HEPES, 150 mM NaCl, 1mM EDTA) was added to the DNA, and incubated for 90 min with 5 μg of S9.6 antibody prebound to magnetic protein A beads. Beads were then washed and the DNA eluted as described above. %RNA–DNA hybrid amounts were quantified using Sybr-Green based quantitative PCRs on DNA samples from DRIP and Input DNA. Q-PCR reactions were performed on ViiA7 Real-Time PCR System (ABI) under standard thermal cycling conditions for 35 cycles. Results were analyzed with ViiA7 software (ABI). For each sample, fold enrichment over Input was calculated by the following equation: Fold enrichment over input = 2^(Ct^Input^ - Log(dilution factor,2) - Ct^DRIP^)

Primer sequences used in this study were adapted from ^32,47,94^ and are listed in Table S4. A mock IP with empty beads was done for each sample to control for the non-specific binding and enrichment was calculated in the same way as the DRIP sample.

### Statistical analysis of Sae2-ChIP-transcription overlap

For each sample, overlap between peaks identified from ChIP-seq data and the top 10% of highly transcribed genes in yeast (Supp. Table 2) ^50–52^ was assessed using bedtools ^95^ and the fraction of peaks for which an overlap was detected was recorded. Also for each sample a background null distribution of overlap rates was estimated by repeatedly sampling a random set of coding sequences equal in number to those in the top 10% list from the Saccharomyces cerevisiae genome (equivalent to selection of top ORFs after random permutation) and then locating overlaps using bedtools. By comparing the overlap fraction to the estimated null distribution, we constructed 99% confidence intervals for the permutation test p-values ^96^ of the null hypothesis that the overlap between ChIP-seq peaks and the top 10% of highly transcribed genes was equal to the rate at which the peaks overlapped randomly chosen coding sequences.

### Mammalian cell culture

Human U2OS, HEK-293T, and MCF7 cells were grown and maintained in tetracycline-free DMEM containing 10% FBS media in a humidified 37°C incubator in the presence of 5% CO_2_.

### Mammalian expression constructs

Invitrogen™ Gateway™ pENTR223 donor vector for hSenataxin(Δ1-1850) was obtained from DNASU (#HsCD00505781), and cloned into pcDNA5/FRT/TO (ThermoFisher) vector to make pTP3531. shRNA CtIP was custom-made by Cellecta (pRSITEP--U6Tet-(sh)-EF1-TetRep-2A-shRNA-CtIP) and includes the shRNA expression cassette 5’- GAGCAGACCTTTCTTAGTATAGTTAATATTCATAGCTATACTGAGAAAGG TCTGCTCTTTT-3 ’. Dox-inducible elements were removed from pRSITEP--U6Tet-(sh)- EF1-TetRep-2A-shRNA-CtIP shRNA CtIP(-Dox) (pTP3914) using Q5® Site-Directed Mutagenesis (NEB, #E0554S). The N-terminal fusion of eGFP with CtIP (eGFP-CtIP) ORF was amplified from pC1-eGFP-CtIP (a generous gift from Steven Jackson), and cloned into pcDNA5/FRT/TO to make pTP3146. The A206K mutation in eGFP was made to prevent eGFP dimerization, and shRNA resistance mutations introduced to generate pTP3148. pcDNA5-flag-bio-CtIP(wt) (containing N-terminal Flag® and biotinylation signal sequences) (pTP3663) was made by Q5® Site-Directed Mutagenesis to remove eGFP, and insert Flag and biotinylation signal sequences. pcDNA5-flag-bio-CtIP(N289A/H290A) (pTP3665) was made using QuikChange II Site-Directed Mutagenesis (Agilent Technologies). shRNA SETX1 (pRSITEP--U6Tet-(sh)-EF1- TetRep-2A-HYGRO) (pTP3677) and shRNA XPG (pRSITEP--U6Tet-(sh)-EF1-TetRep-2A-HYGRO) (pTP3703) were made from the pRSITEP--U6Tet-(sh)-EF1-TetRep-2A-shRNA-CtIP construct (Celecta) using Q5® Site-Directed Mutagenesis, with the shRNA cassettes 5’- GCCAGATCGTATACAATTATAGTTAATATTCATAGCTATAATTGTATACGATCTGGCT TTT-3 ’ and 5’- GAACGCACCTGCTGCTGTAGAGTTAATATTCATAGCTCTACAGCAGCAGGTGCGTT CTTTT-3 ’, respectively. pcDNA5-RNaseHI-D10R-E48R-NLS-mCherry (RNaseHI mutant with C-terminal NLS and mCherry tags, respectively) was cloned using pcDNA5/FRO/TO and pICE-RNaseHI-D10R-E48R-NLS-mCherry (a gift from Patrick Calsou, Addgene # 60364 ^97^), pcDNA5-NLS-mCherry (pTP3494) was cloned using pcDNA5/FRO/TO and pICE-NLS-mCherry (a gift from Patrick Calsou, Addgene, # 60367)). pLenti-PGK-RNaseHI-D10R-E48R-NLS-mCherry (pTP3660) was cloned using pLenti PGK GFP Blast (w510-5) which was a gift from Eric Campeau & Paul Kaufman (Addgene plasmid #19069 ^98^) and the RNaseHI-D10R-E48R-NLS-mCherry fragment from pICE-RNaseHI-D10R-E48R-NLS-mCherry. pBluescript-Sγ3×12 was a generous gift from Lieber lab (pTW-121), pcDNA5/FRT/TO-Sγ3×12 (pTP4195) was cloned using pcDNA5/FRT/TO and the Sγ3 region from pTW-121. pcDNA5/FRT/TO-hPCF11 (pTP4195) was cloned using pcDNA5/FRT/TO and the hPCF11 ORF containing plasmid, obtained from Kazusa (ORK06290). All constructs and mutations were confirmed by DNA sequencing. Details of plasmid construction available upon request.

### Clonogenic survival assays

U2OS cells were harvested by trypsinization (0.25% trypsin; Life Technologies) and counted using Scepter (Millipore Sigma). 1,500 cells were seeded per 10 cm cell culture dish, and were allowed to adhere to the bottom of the plate for 36 hrs. For experiments with tet-inducible CtIP shRNA, all cells were exposed to doxycycline (1 µg/mL) during seeding and gene depletion and/or over-expression and throughout the experimental course. After 36 hours cells were treated CPT for 1 hour, the CPT-containing media was replaced with fresh media, and cell recovery was allowed for 10 days. Colonies were stained with crystal violet (0.05% in 20% ethanol), destained with water, and counted using a scanner and Image J.

### RNaseH^*D10R-E48R*^-mCherry Laser Micro-irradiation

U2OS cells were seeded in glass bottom petri dishes (35×10 mm, 22 mm glass, WillCo-dish®, HBST-3522), and grown in DMEM / 10% FBS media in the presence of 1 µg/mL doxycycline. After 36 hrs, media was replaced with media containing 10 µM BrdU. After 36 hours, laser micro-irradiation was performed with an inverted confocal microscope (FV1000; Olympus) equipped with a CO_2_ module and a 37°C heating chamber. A preselected spot within the nucleus was microirradiated with 20 iterations of a 405-nm laser with 100% power to generate localized DNA damage. Then, time-lapse images were acquired using a red laser at 1 min time intervals for 10 min. The fluorescence intensity of mCherry signal at the laser microirradiated sites was measured using the microscope’s software. Data collected from >10 cells were normalized to their initial intensity and plotted against time.

### RNaseH^*D10R-E48R*^-mCherry FACS

U2OS cells were grown in DMEM/10% FBS media in the presence of 1 µg/mL doxicyclin in 10 cm dishes. After 3 days, the cells were harvested by trypsinization, rinsed in 5 mL cold PBS with Ca^2+^ (0.9 mM) and Mg^2+^ (0.5 mM), and centrifuged at 1000 g for 3 min. Unbound RNaseH^D10R-E48R^-mCherry was extracted with 1 mL Triton X-100 buffer (0.5% Triton X-100, 20 mM Hepes-KOH (pH 7.9), 50 mM NaCl, 3 mM MgCl_2_, 300 mM Sucrose) at 4°C for exactly 2 min and at 1300 g for 3 min. Cells were then rinsed twice in PBS at RT and fixed in 3.7% paraformaldehyde in PBS for 10 min at RT, rinsed twice in PBS at RT again, and stained with FxCycleTM FarRed stain (200 nM in PBS with 100 µg/mL RNaseA). Cells were kept in the staining solution at 4°C protected from light. The samples were analyzed in a flow cytometer without washing, using 633/5 nm excitation and emission collected in a 660/20 band pass or equivalent.

### S9.6 immunostaining

U2OS cells were seeded into 8-well Nunc™ Lab-Tek™ II Chamber Slides™ (Nalge Nunc International, #154534) 48 hrs before experiments. Prior to immunostaining, cells were washed with PBS, and preextracted with incubation in CSK buffer (10 mM PIPES, pH 7.0, 100 mM NaCl, 300 mM sucrose, and 3 mM MgCl_2_, 0.7% Triton X-100) twice for 3 min at room temperature. After preextraction, cells were washed with PBS and fixed with 2% paraformaldehyde. Cells were then permeabilized for 5 min with PBS/0.2% Triton X-100, washed with PBS, and blocked with PBS/0.1% Tween 20 (PBS-T) containing 5% BSA. For immunostaining cells were incubated with primary antibodies (S9.6 and nucleolin) in PBS/5% BSA overnight, then washed with PBS-T and incubated with appropriate secondary antibodies coupled to Alexa Fluor 488 or 594 fluorophores (Life Technologies) in PBS-T/5% BSA. After washes in PBS-T and PBS, coverslips were incubated 30 min with 2 µg/ml DAPI in PBS. After washes in PBS, coverslips were rinsed with water and mounted on glass slides using ProLong Gold (Life Technologies).

### DNA-RNA ImmunoPrecipitation (DRIP)

U2OS cells (one 150 mm dish per biological replicate) were harvested by trypsinization (0.25% trypsin; Life Technologies) and pelleted at 1,000 RCF for 5 min in 15 mL Falcon tubes. Cell pellets were washed with PBS and divided for RNA, DRIP or LM-PCR harvests. Cell pellets for DRIP were resuspended in 5 mL of PBS supplemented with 0.5% SDS, and digested with 2 mg of Proteinase K (GoldBio) at 37°C overnight. Cell lysates were then extracted once with 1 volume of equilibrated phenol pH 8 and twice with 1 volume of chloroform. DNA was precipitated with 1 volume of isopropanol, and spun down at 6,500 RCF for 15 min. The DNA pellet was transferred to a 1.7 mL eppendorf tube, washed with 1 mL of 70% ethanol, and rehydrated in 0.1x TE (1 mM Tris-HCl pH 8.0, 0.1 mM EDTA). Nucleic acids were digested using a restriction enzyme cocktail (20 units each of EcoRI, HindIII, BsrGI, XbaI) (New England Biolabs) overnight at 37°C in 1x NEBuffer 2.1. Digests DNA concentration was measured using Qubit dsDNA HS kit (Thermofisher). 10 µg of digested nucleic acids were diluted in 1 mL final DRIP buffer (10 mM sodium phosphate, 140 mM sodium chloride, 0.05% Triton X-100) and 100 µg of S9.6 antibody (purified from ATCC HB-8730) and incubated at 4°C overnight. This and all wash steps were performed on a rotisserie mixer. 30 µL of Pierce Protein A/G Magnetic Beads (Fisher Scientific, 88803) was added and incubated for additional 2 hrs, followed by washing three times with 1 mL of 1x IP buffer for 10 min at room temperature with constant rotation. After the final wash, the agarose slurry was resuspended in 100 µL of TE + 0.5% SDS 1 mg of Proteinase K for > 1 hr at 37°C. 10 µL of 7.5 M Ammonium Acetate, 1 µg of glycogen, and 400 µL of 100% ice-cold Ethanol were added to the digested DRIP samples, and kept at −20°C for at least 2 hrs (to overnight) to precipitate the immuno-precipitated material. The pellet was precipitated by centrifugation in a microcentrifuge at maximum speed for 30 min at 4°C, washed with cold 70% ethanol, air-dried, and resuspended in 100 µL of 0.1x TE. We used 10 µL reactions with PowerUp® sybr green master mix (Applied Biosystems) for qPCR amplification of genomic loci (see Table S4). Reactions were incubated with the following program on a Viia 7 System (Life Technologies): 50°C 2 minutes, 95°C 10 minutes, 40 cycles of 95°C 15 seconds, 64°C 1 minute, followed by a melt curve: 95°C 15 seconds, 60°C 1 minute, 0.05°C/second to 95°C 15 seconds. For each DRIP sample, linear range of amplification was identified by testing a wide range of dilutions. Fold enrichment for a given locus was calculated using the 2-I:) I:) CT method ^99^, and then normalizing the samples to the measurements of the wild-type results.

### RT-qPCR

U2OS cells (one 150 mm dish per biological replicate) were harvested by trypsinization (0.25% trypsin; Life Technologies) and pelleted at 1,000 RCF for 5 min in 15 mL Falcon tubes. Cell pellets were washed with PBS (Life Technologies) and divided for RNA, DRIP or LM-PCR harvests. RNA was either purified and retro-transcribed using RNA purification (Qiagen) and SuperScript® IV Reverse Transcriptase (Thermo Fisher Scientific, #8090050) kits, respectively, or using Fast Cells-to-CT kit (Thermo Fisher Scientific, #4399003). qPCR was done using the same settings as those for DRIP-qPCR and LM-PCR methods, and GAPDH used as a reference gene.

### mRNA-seq

U2OS cells (one 150 mm dish per biological replicate) were harvested by trypsinization and pelleted at 1,000 RCF for 5 min in 15 mL Falcon tubes. mRNA was purified using Qiagen RNA purification kit, and mRNA was isolated using AMPure XP kit (Beckman Coulter, #A63880). mRNA-seq libraries were prepared with the NEBNext® Multiplex Small RNA Library Prep Set for Illumina® (#E7300S) according to manufacturer instructions. The library sequencing and analysis were done at New York Genome Center.

### Statistical analysis of mRNA-seq DRIP data overlap

For each statistical comparison, overlap between the top 100 genes as ranked by DESeq differential expression p-value and DRIPc-seq peaks from GEO dataset GSE70189 (Sanz 2016) was assessed using bedtools ^95^. For each comparison, we calculated the percentage of the top 100 genes for which such an overlap was found. These percentages were compared to a null distribution for overlap rates estimated using a permutation testing approach (Ernst 2004) in which we repeatedly selected 100 genes at random from the list of all genes tested by DESeq (equivalent to selection of top 100 genes after randomly permuting DESeq p-values) and applied bedtools to calculate the DRIPc-seq overlap rate in the same manner. Using this approach we estimated 99% confidence intervals for the permutation test p-values of the null hypothesis that the genes showing the most evidence of differential expression overlapped the peak regions at the same rate as randomly selected genes.

### Ligation-mediated PCR (LM-PCR)

U2OS cells (one 150 mm dish per biological replicate) were harvested by trypsinization and pelleted at 1,000 RCF for 5 min in 15 mL Falcon tubes. Cell pellets were washed with PBS and divided for RNA, DRIP or LM-PCR harvests. Genomic DNA (gDNA) was purified using genomic DNA preparation kit (Zymo Research Quick-gDNA™ MiniPrep - Capped column, Genesee Scientific, 11-317AC) and gDNA concentration was determined using Nanodrop. 50 µL primer extension mix contained: 5 µL of 10X polymerase buffer (NEB, supplied with Deepvent(-Exo) enzme), 4 µL MgSO_4_ (100 mM), 1 µL of Deepvent(-Exo) (NEB), 1 µL NTP mix (0.5 mM each final), 0.5 µL biotinylated primers (stock concentration 100 µM), and 1 µg of gDNA. Primer extension was done in a thermocycler in one round of primer extension: 15 min at 95°C, 30 sec at 60°C, 5 min @ 72°C. Control solution was made by dilution of 1 µg genomic DNA in 0.1x TE. Primer extension products were ligated to the phosphorylated asymmetric adaptor duplex overnight (oligonucleotides were phosphorylated with T4 polynucleotide kinase at 37°C for 3-4 hrs, purified with a nucleotide removal kit (Qiagen), and annealed with boiling and slow cooling in the presence of 0.1 M NaCl). Ligation reactions were mixed with 30 µL of KilobaseBinder (Invitrogen) magnetic beads prepared according to the manufacturer’s protocol, total volume was adjusted to 100 µL, and incubated with genomic DNA samples overnight. Washes were performed on a magnetic stand: 3x 10 min washing with wash buffer (50 mM Tris, pH 8, 0.1% (wt/vol) SDS and 150 mM NaCl), then 10 min washing with 0.1x TE. After the 0.1x TE wash, the beads were resuspended in 100 µL of 0.1x TE and 10 µL used for nested PCR. Nested PCR reaction contained: 5 µL of 10X polymerase buffer (NEB, supplied with Deepvent(-Exo) enzme), 4 µL MgSO_4_ (100 mM), 1 µL of Deepvent(-Exo), 1 µL NTP mix (0.5 mM each final), 1 µL each nested primers (stock concentration 100 µM), and 10 µL of beads. Nested PCR was done in a thermocycle with the following amplification steps: 1) one step of total denaturation: 15 min at 95°C; 2) 15 steps of amplification: 30 sec at 95°C 30 sec at 60°C, 5 min @ 72°C; and 3) one step of extension: 5 min @ 72°C. Nested reactions were diluted 50-fold in 0.1x TE, and serial dilutions were prepared to determine linear range of amplification. Fold enrichment for a given locus was calculated using the comparative Ct method, and then normalizing the samples to the measurements of the wild-type results.

### Comet assay

U2OS cells were grown in DMEM/10% FBS media in the presence of 1 µg/mL doxicyclin in 6-well plates at a very sparse seeding density. After 3 days, the cells were treated with DNA damaging agents, harvested by trypsinization, and rinsed in 1 mL cold PBS. Olive moments of damaged DNA were measured using OxiSelect™ Comet Assay Kit (3-Well Slides) (Cellbiolabs, #STA-350).

### R-loops in vitro

The eGFP ORF was cloned into the topo-pSC-A-amp/kan vector (Agilent Technologies) to make pTP3823, which was purified using the alkaline CsCl method. The negative supercoiling of the purified plasmid was removed by the Nt.BspQI nickase treatment and the relaxed plasmid was used as a template for the T7 RNA polymerase-mediated transcription reactions. Briefly, 1 µg of DNA was transcribed with 20 units of T7 polymerase (T7 polymerase kit, Promega) in 10 µL total reaction volume in the presence of radioactive alpha UTP (0.5 mM ATP, 0.5 mM CTP, 0.5 mM GTP, 0.1 mM UTP, and 10 µCi ^32^P apha-UTP) for one hour at 37 °C, then 10 µL of various amounts of the S1 nuclease (0, 0.01, 0.05, and 2.5 units, respectively) (Thermo Fisher Scientific, #EN0321) in the corresponding 2X nuclease buffer (total volume 20 µL) was added to the transcription reactions for two additional hrs. After a total of three hours, reactions were stopped by adding 2 µL of 0.5 M EDTA, 2 µL of 5 M NaCl, and 1 µg RNaseA for 15 min, then deproteinated with 1 µg of Proteinase K in 0.5 % SDS. The reaction products were ethanol-precipitated (1/10 volume of NH_4_OAc, 4 volumes of 100% EtOH, kept at −20 °C overnight), treated with 50 units of RNase H in the respective buffer where applicable, and resolved on a 1% agarose gel in 1X TBE for 2.5 hours at 120 V. Gels were stained with EtBr for 30 min, and visualized on UV platen, then dried and exposed to phospho-screens. Phospho-screens were visualized using Typhoon FLA 9500.

## Acknowledgements

Work in the T.T.P. laboratory is supported in part by the Cancer Prevention and Research Institute of Texas (RP160667). The K.M.M. laboratory is supported by the NIH National Cancer Institute (R01CA198279 and RO1CA201268) and the American Cancer Society (RSG-16-042-01-DMC). The work of Justin Leung is supported by the NIH National Cancer Institute (K22CA204354). We thank Steve Hanes, Nicholas Proudfoot, Jeff Corden, John Petrini, Hannah Klein, Lorraine Symington, Jim Haber, Michael Lieber, Steve Jackson, Patrick Calsou, Eric Campeau, and Paul Kaufman for reagents including yeast strains and plasmids.

## Author contributions

Q.F., S.A., Y.Y., and T.T.P. performed experiments in budding yeast; N.M. and J.L.performed experiments in human cells; K.M.M., T.T.P. and all the authors contributed to the writing of the manuscript.

## Competing Interests

The authors declare no competing interests.

## Materials & Correspondence

Correspondence and requests for materials should be addressed to Tanya Paull

## References

1. Aparicio, T., Baer, R. & Gautier, J. DNA double-strand break repair pathway choice and cancer. DNA Repair Amst 19, 169–75 (2014).

2. Ceccaldi, R., Rondinelli, B. & D’Andrea, A. D. Repair Pathway Choices and Consequences at the Double-Strand Break. Trends Cell Biol. 26, 52–64 (2016).

3. Symington, L. S. Mechanism and regulation of DNA end resection in eukaryotes. Crit Rev Biochem Mol Biol 51, 195–212 (2016).

4. Symington, L. S. & Gautier, J. Double-strand break end resection and repair pathway choice. Annu Rev Genet 45, 247–71 (2011).

5. Cannavo, E. & Cejka, P. Sae2 promotes dsDNA endonuclease activity within Mre11-Rad50-Xrs2 to resect DNA breaks. Nature 514, 122–5 (2014).

6. Reginato, G., Cannavo, E. & Cejka, P. Physiological protein blocks direct the Mre11-Rad50-Xrs2 and Sae2 nuclease complex to initiate DNA end resection. Genes Dev 31, 2325–2330 (2017).

7. Wang, W., Daley, J. M., Kwon, Y., Krasner, D. S. & Sung, P. Plasticity of the Mre11-Rad50-Xrs2-Sae2 nuclease ensemble in the processing of DNA-bound obstacles. Genes Dev 31, 2331–2336 (2017).

8. Cejka, P. et al. DNA end resection by Dna2-Sgs1-RPA and its stimulation by Top3-Rmi1 and Mre11-Rad50-Xrs2. Nature 467, 112–6 (2010).

9. Myler, L. R. et al. Single-molecule imaging reveals the mechanism of Exo1 regulation by single-stranded DNA binding proteins. Proc Natl Acad Sci U A 113, E1170–9 (2016).

10. Nicolette, M. L. et al. Mre11-Rad50-Xrs2 and Sae2 promote 5’ strand resection of DNA double-strand breaks. Nat Struct Mol Biol 17, 1478–85 (2010).

11. Niu, H. et al. Mechanism of the ATP-dependent DNA end-resection machinery from Saccharomyces cerevisiae. Nature 467, 108–11 (2010).

12. Shim, E. Y. et al. Saccharomyces cerevisiae Mre11/Rad50/Xrs2 and Ku proteins regulate association of Exo1 and Dna2 with DNA breaks. EMBO J 29, 3370–3380 (2010).

13. Chanut, P., Britton, S., Coates, J., Jackson, S. P. & Calsou, P. Coordinated nuclease activities counteract Ku at single-ended DNA double-strand breaks. Nat Commun 7, 12889 (2016).

14. Makharashvili, N. et al. Catalytic and Noncatalytic Roles of the CtIP Endonuclease in Double-Strand Break End Resection. Mol. Cell 54, 1022–33 (2014).

15. Wang, H. et al. An end resection-independent CtIP endonuclease activity is required for maintaining genome stability at common fragile sites and inverted repeats. Mol. Cell (2014).

16. Deng, C., Brown, J. A., You, D. & Brown, J. M. Multiple endonucleases function to repair covalent topoisomerase I complexes in Saccharomyces cerevisiae. Genetics 170, 591–600 (2005).

17. Sartori, A. A. et al. Human CtIP promotes DNA end resection. Nature 450, 509–14 (2007).

18. Pommier, Y. Topoisomerase I inhibitors: camptothecins and beyond. Nat. Rev. Cancer 6, 789–802 (2006).

19. Liu, L. F. & Wang, J. C. Supercoiling of the DNA template during transcription. Proc. Natl. Acad. Sci. U. S. A. 84, 7024–7027 (1987).

20. Pommier, Y., Sun, Y., Huang, S. N. & Nitiss, J. L. Roles of eukaryotic topoisomerases in transcription, replication and genomic stability. Nat. Rev. Mol. Cell Biol. 17, 703–721 (2016).

21. Zhang, H., Wang, J. C. & Liu, L. F. Involvement of DNA topoisomerase I in transcription of human ribosomal RNA genes. Proc. Natl. Acad. Sci. U. S. A. 85, 1060–1064 (1988).

22. McKee, A. H. & Kleckner, N. A general method for identifying recessive diploid-specific mutations in Saccharomyces cerevisiae, its application to the isolation of mutants blocked at intermediate stages of meiotic prophase and characterization of a new gene SAE2. Genetics 146, 797–816 (1997).

23. Neale, M. J., Pan, J. & Keeney, S. Endonucleolytic processing of covalent protein-linked DNA double-strand breaks. Nature 436, 1053–7 (2005).

24. Prinz, S., Amon, A. & Klein, F. Isolation of COM1, a new gene required to complete meiotic double-strand break-induced recombination in Saccharomyces cerevisiae. Genetics 146, 781–95 (1997).

25. Sollier, J. & Cimprich, K. A. Breaking bad: R-loops and genome integrity. Trends Cell Biol. 25, 514–522 (2015).

26. Santos-Pereira, J. M. & Aguilera, A. R loops: new modulators of genome dynamics and function. Nat. Rev. Genet. 16, 583–597 (2015).

27. Costantino, L. & Koshland, D. The Yin and Yang of R-loop biology. Curr. Opin. Cell Biol. 34, 39–45 (2015).

28. Hamperl, S., Bocek, M. J., Saldivar, J. C., Swigut, T. & Cimprich, K. A. Transcription-Replication Conflict Orientation Modulates R-Loop Levels and Activates Distinct DNA Damage Responses. Cell 170, 774–786.e19 (2017).

29. Finkel, J. S., Chinchilla, K., Ursic, D. & Culbertson, M. R. Sen1p performs two genetically separable functions in transcription and processing of U5 small nuclear RNA in Saccharomyces cerevisiae. Genetics 184, 107–18 (2010).

30. Mischo, H. E. et al. Yeast Sen1 helicase protects the genome from transcription-associated instability. Mol. Cell 41, 21–32 (2011).

31. Steinmetz, E. J. et al. Genome-wide distribution of yeast RNA polymerase II and its control by Sen1 helicase. Mol. Cell 24, 735–46 (2006).

32. Alzu, A. et al. Senataxin associates with replication forks to protect fork integrity across RNA-polymerase-II-transcribed genes. Cell 151, 835–46 (2012).

33. Cohen, S. et al. Senataxin resolves RNA:DNA hybrids forming at DNA double-strand breaks to prevent translocations. Nat. Commun. 9, 533 (2018).

34. Skourti-Stathaki, K., Proudfoot, N. J. & Gromak, N. Human senataxin resolves RNA/DNA hybrids formed at transcriptional pause sites to promote Xrn2-dependent termination. Mol. Cell 42, 794–805 (2011).

35. Yüce, Ö. & West, S. C. Senataxin, defective in the neurodegenerative disorder ataxia with oculomotor apraxia 2, lies at the interface of transcription and the DNA damage response. Mol. Cell. Biol. 33, 406–417 (2013).

36. Groh, M., Albulescu, L. O., Cristini, A. & Gromak, N. Senataxin: Genome Guardian at the Interface of Transcription and Neurodegeneration. J. Mol. Biol. 429, 3181–3195 (2017).

37. Porrua, O. & Libri, D. Transcription termination and the control of the transcriptome: why, where and how to stop. Nat. Rev. Mol. Cell Biol. 16, 190–202 (2015).

38. Moreau, S., Ferguson, J. R. & Symington, L. S. The nuclease activity of Mre11 is required for meiosis but not for mating type switching, end joining, or telomere maintenance. Mol Cell Biol 19, 556–66 (1999).

39. Chinchilla, K. et al. Interactions of Sen1, Nrd1, and Nab3 with Multiple Phosphorylated Forms of the Rpb1 C-Terminal Domain in Saccharomyces cerevisiae. Eukaryot. Cell 11, 417–429 (2012).

40. Vaze, M. B. et al. Recovery from checkpoint-mediated arrest after repair of a double-strand break requires Srs2 helicase. Mol Cell 10, 373–85 (2002).

41. Clerici, M., Mantiero, D., Lucchini, G. & Longhese, M. P. The Saccharomyces cerevisiae Sae2 protein promotes resection and bridging of double strand break ends. J Biol Chem 280, 38631–8 (2005).

42. DeMarini, D. J., Winey, M., Ursic, D., Webb, F. & Culbertson, M. R. SEN1, a positive effector of tRNA-splicing endonuclease in Saccharomyces cerevisiae. Mol. Cell. Biol. 12, 2154–2164 (1992).

43. Arudchandran, A. et al. The absence of ribonuclease H1 or H2 alters the sensitivity of Saccharomyces cerevisiae to hydroxyurea, caffeine and ethyl methanesulphonate: implications for roles of RNases H in DNA replication and repair. Genes Cells Devoted Mol. Cell. Mech. 5, 789–802 (2000).

44. Lazzaro, F. et al. RNase H and Postreplication Repair Protect Cells from Ribonucleotides Incorporated in DNA. Mol. Cell 45, 99–110 (2012).

45. Zimmer, A. D. & Koshland, D. Differential roles of the RNases H in preventing chromosome instability. Proc. Natl. Acad. Sci. 113, 12220–12225 (2016).

46. Birse, C. E., Minvielle-Sebastia, L., Lee, B. A., Keller, W. & Proudfoot, N. J. Coupling termination of transcription to messenger RNA maturation in yeast. Science 280, 298–301 (1998).

47. Grzechnik, P., Gdula, M. R. & Proudfoot, N. J. Pcf11 orchestrates transcription termination pathways in yeast. Genes Dev 29, 849–61 (2015).

48. Jimenez, A., Tipper, D. J. & Davies, J. Mode of action of thiolutin, an inhibitor of macromolecular synthesis in Saccharomyces cerevisiae. Antimicrob Agents Chemother 3, 729–38 (1973).

49. Zhang, Y. et al. Model-based analysis of ChIP-Seq (MACS). Genome Biol. 9, R137 (2008).

50. Miura, F. et al. Absolute quantification of the budding yeast transcriptome by means of competitive PCR between genomic and complementary DNAs. BMC Genomics 9, 574 (2008).

51. Nagalakshmi, U. et al. The Transcriptional Landscape of the Yeast Genome Defined by RNA Sequencing. Science 320, 1344–1349 (2008).

52. Pelechano, V., Chávez, S. & Pérez-Ortín, J. E. A Complete Set of Nascent Transcription Rates for Yeast Genes. PLoS ONE 5, e15442 (2010).

53. Boguslawski, S. J. et al. Characterization of monoclonal antibody to DNA.RNA and its application to immunodetection of hybrids. J. Immunol. Methods 89, 123–130 (1986).

54. Schaughency, P., Merran, J. & Corden, J. L. Genome-Wide Mapping of Yeast RNA Polymerase II Termination. PLoS Genet. 10, e1004632 (2014).

55. Makharashvili, N. & Paull, T. T. CtIP: A DNA damage response protein at the intersection of DNA metabolism. DNA Repair Amst 32, 75–81 (2015).

56. Huertas, P. & Jackson, S. P. Human CtIP mediates cell cycle control of DNA end resection and double strand break repair. J Biol Chem 284, 9558–65 (2009).

57. Nakamura, K. et al. Collaborative action of Brca1 and CtIP in elimination of covalent modifications from double-strand breaks to facilitate subsequent break repair. PLoS Genet 6, e1000828 (2010).

58. Bhatia, V. et al. BRCA2 prevents R-loop accumulation and associates with TREX-2 mRNA export factor PCID2. Nature 511, 362–365 (2014).

59. Lavin, M. F., Yeo, A. J. & Becherel, O. J. Senataxin protects the genome: Implications for neurodegeneration and other abnormalities. Rare Dis. Austin Tex 1, e25230 (2013).

60. Suraweera, A. et al. Senataxin, defective in ataxia oculomotor apraxia type 2, is involved in the defense against oxidative DNA damage. J. Cell Biol. 177, 969–979 (2007).

61. Fagbemi, A. F., Orelli, B. & Schärer, O. D. Regulation of endonuclease activity in human nucleotide excision repair. DNA Repair 10, 722–729 (2011).

62. Sollier, J. et al. Transcription-coupled nucleotide excision repair factors promote R-loop-induced genome instability. Mol Cell 56, 777–85 (2014).

63. Chen, P. L. et al. Inactivation of CtIP leads to early embryonic lethality mediated by G1 restraint and to tumorigenesis by haploid insufficiency. Mol Cell Biol 25, 3535–42 (2005).

64. Polato, F. et al. CtIP-mediated resection is essential for viability and can operate independently of BRCA1. J. Exp. Med. 211, 1027–1036 (2014).

65. García-Rubio, M., Barroso, S. I. & Aguilera, A. Detection of DNA-RNA Hybrids In Vivo. in Genome Instability (eds. Muzi-Falconi, M. & Brown, G. W.) 1672, 347–361 (Springer New York, 2018).

66. Schaeper, U., Subramanian, T., Lim, L., Boyd, J. M. & Chinnadurai, G. Interaction between a cellular protein that binds to the C-terminal region of adenovirus E1A (CtBP) and a novel cellular protein is disrupted by E1A through a conserved PLDLS motif. J. Biol. Chem. 273, 8549–8552 (1998).

67. Hatchi, E. et al. BRCA1 Recruitment to Transcriptional Pause Sites Is Required for R-Loop-Driven DNA Damage Repair. Mol. Cell 57, 636–647 (2015).

68. Monteiro, A. N., August, A. & Hanafusa, H. Evidence for a transcriptional activation function of BRCA1 C-terminal region. Proc. Natl. Acad. Sci. U. S. A. 93, 13595–13599 (1996).

69. Scully, R. et al. BRCA1 is a component of the RNA polymerase II holoenzyme. Proc Natl Acad Sci U A 94, 5605–10 (1997).

70. Takaoka, M. & Miki, Y. BRCA1 gene: function and deficiency. Int. J. Clin. Oncol. 23, 36–44 (2018).

71. Yu, X., Wu, L. C., Bowcock, A. M., Aronheim, A. & Baer, R. The C-terminal (BRCT) domains of BRCA1 interact in vivo with CtIP, a protein implicated in the CtBP pathway of transcriptional repression. J Biol Chem 273, 25388–92 (1998).

72. Ginno, P. A., Lott, P. L., Christensen, H. C., Korf, I. & Chédin, F. R-loop formation is a distinctive characteristic of unmethylated human CpG island promoters. Mol. Cell 45, 814–825 (2012).

73. Huang, F.-T., Yu, K., Hsieh, C.-L. & Lieber, M. R. Downstream boundary of chromosomal R-loops at murine switch regions: Implications for the mechanism of class switch recombination. Proc. Natl. Acad. Sci. 103, 5030–5035 (2006).

74. Bhatia, V. et al. BRCA2 prevents R-loop accumulation and associates with TREX-2 mRNA export factor PCID2. Nature 511, 362–365 (2014).

75. Skourti-Stathaki, K., Kamieniarz-Gdula, K. & Proudfoot, N. J. R-loops induce repressive chromatin marks over mammalian gene terminators. Nature 516, 436–439 (2014).

76. Baranello, L. et al. RNA Polymerase II Regulates Topoisomerase 1 Activity to Favor Efficient Transcription. Cell 165, 357–371 (2016).

77. Khobta, A. et al. Early effects of topoisomerase I inhibition on RNA polymerase II along transcribed genes in human cells. J. Mol. Biol. 357, 127–138 (2006).

78. Merino, A., Madden, K. R., Lane, W. S., Champoux, J. J. & Reinberg, D. DNA topoisomerase I is involved in both repression and activation of transcription. Nature 365, 227–232 (1993).

79. Arora, S. et al. Genetic Separation of Sae2 Nuclease Activity from Mre11 Nuclease Functions in Budding Yeast. Mol Cell Biol 37, (2017).

80. Roy, D., Zhang, Z., Lu, Z., Hsieh, C.-L. & Lieber, M. R. Competition between the RNA Transcript and the Nontemplate DNA Strand during R-Loop Formation In Vitro: a Nick Can Serve as a Strong R-Loop Initiation Site. Mol. Cell. Biol. 30, 146–159 (2010).

81. Martin-Tumasz, S. & Brow, D. A. Saccharomyces cerevisiae Sen1 Helicase Domain Exhibits 5’-to 3’-Helicase Activity with a Preference for Translocation on DNA Rather than RNA. J. Biol. Chem. 290, 22880–22889 (2015).

82. Wahba, L., Costantino, L., Tan, F. J., Zimmer, A. & Koshland, D. S1-DRIP-seq identifies high expression and polyA tracts as major contributors to R-loop formation. Genes Dev. 30, 1327–1338 (2016).

83. Chinnadurai, G. CtIP, a candidate tumor susceptibility gene is a team player with luminaries. Biochim Biophys Acta 1765, 67–73 (2006).

84. Hill, S. J. et al. Systematic screening reveals a role for BRCA1 in the response to transcription-associated DNA damage. Genes Dev. 28, 1957–1975 (2014).

85. Trego, K. S. et al. Non-catalytic Roles for XPG with BRCA1 and BRCA2 in Homologous Recombination and Genome Stability. Mol Cell 61, 535–46 (2016).

86. Barton, O. et al. Polo-like kinase 3 regulates CtIP during DNA double-strand break repair in G1. J Cell Biol 206, 877–94 (2014).

87. Biehs, R. et al. DNA Double-Strand Break Resection Occurs during Non-homologous End Joining in G1 but Is Distinct from Resection during Homologous Recombination. Mol Cell 65, 671–684 e5 (2017).

88. Helmink, B. A. et al. H2AX prevents CtIP-mediated DNA end resection and aberrant repair in G1-phase lymphocytes. Nature 469, 245–9 (2011).

89. Quennet, V., Beucher, A., Barton, O., Takeda, S. & Lobrich, M. CtIP and MRN promote non-homologous end-joining of etoposide-induced DNA double-strand breaks in G1. Nucleic Acids Res 39, 2144–52 (2011).

90. Yun, M. H. & Hiom, K. CtIP-BRCA1 modulates the choice of DNA double-strand-break repair pathway throughout the cell cycle. Nature 459, 460–3 (2009).

91. Adams, A., Gottschling, D. E. & Kaiser, C. Methods in Yeast Genetics. (Cold Spring Harbor Laboratory Press, 1997).

92. Christianson, T. W., Sikorski, R. S., Dante, M., Shero, J. H. & Hieter, P. Multifunctional yeast high-copy-number shuttle vectors. Gene 110, 119–22 (1992).

93. Fu, Q. et al. Phosphorylation-regulated transitions in an oligomeric state control the activity of the sae2 DNA repair enzyme. Mol Cell Biol 34, 778–93 (2014).

94. El Hage, A., Webb, S., Kerr, A. & Tollervey, D. Genome-wide distribution of RNA-DNA hybrids identifies RNase H targets in tRNA genes, retrotransposons and mitochondria. PLoS Genet. 10, e1004716 (2014).

95. Quinlan, A. R. & Hall, I. M. BEDTools: a flexible suite of utilities for comparing genomic features. Bioinformatics 26, 841–842 (2010).

96. Ernst, M. D. Permutation Methods: A Basis for Exact Inference. Stat. Sci. 19, 676–685 (2004).

97. Britton, S. et al. DNA damage triggers SAF-A and RNA biogenesis factors exclusion from chromatin coupled to R-loops removal. Nucleic Acids Res. 42, 9047–9062 (2014).

98. Campeau, E. et al. A versatile viral system for expression and depletion of proteins in mammalian cells. PloS One 4, e6529 (2009).

99. Schmittgen, T. D. & Livak, K. J. Analyzing real-time PCR data by the comparative CT method. Nat. Protoc. 3, 1101–1108 (2008).

